# Material matters: raw material influences stone tool performance in capuchin monkeys

**DOI:** 10.1101/2024.12.12.628091

**Authors:** Theo D. R. O’Malley, Jonathan S. Reeves, Nora E. Slania, Tiago Falótico, Tomos Proffitt, Ignacio de la Torre, Lydia V. Luncz

## Abstract

Identifying the conditions that facilitate tool use is a central focus in the field of human evolution and animal behavior. Particular interest lies in studying stone tool use by non-human primates, especially when stone hammers are used to open encased food sources. Percussive tool use is a rare and complex technological behavior which was also employed by our early hominin ancestors. It is widely theorized that environmental factors play a role in explaining the presence and absence of stone tool use across primate populations. Research to date has focused heavily on the quality and availability of tool-extractable foods, and how these compare to alternative food sources. There has been limited investigation into whether access to different types of raw tool material for hammerstones and anvils affects the efficiency of tool users when exploiting encased resources. Here, we quantitatively analyzed how percussive tools of different raw materials vary in their performance and durability. Wild capuchins (*Sapajus libidinosus*) in Brazil were provided with stones sourced globally from primate and hominin tool use sites. We measured the reliability with which monkeys could crack nuts when using different raw materials, the number of strikes required to open a nut or obtain a food reward, and how these metrics changed over time. We further reported variations in the durability of different raw materials which directly relates to how long a tool remains usable. Our results showed that differences in nut cracking reliability and efficiency were largely driven by the ability of the tool material to stabilize the nut. Furthermore, there was wide variation in tool durability, particularly for anvils. These results highlight the importance of considering raw materials among the ecological factors that can influence the selective benefits or costs of tool use behaviors.

**HIGHLIGHTS:**

- Raw material influences primate percussive stone tool durability and performance
- Materials differ in their ability to stabilize food sources
- Efficiency of hammer strikes differed across materials
- Differences in material performance did not depend on their capacity to from pits
- Material performance has consequences for food yield and energetic efficiency

## INTRODUCTION

The adoption of stone tools by early hominins laid the foundation for incremental increases in cognitive and technical abilities culminating in a complex technological culture amongst humans (De Beaune, 2004; Langley & Suddendorf, 2022). Stone tool use has since been reported for six non-human primate species (hereafter ‘primates’), including the western chimpanzee (*Pan troglodytes verus*; Boesch & Boesch, 1990), bearded, yellow-breasted, blond, and white-face capuchins (*Sapajus libidinosus*; Falótico & Ottoni, 2016; *S. xanthosternos*; Canale et al., 2009; *S. flavius*; Lima et al., 2024; *Cebus capucinus*; Barrett et al., 2018), and Burmese long-tailed macaques (*Macaca fascicularis aurea*; Malaivijitnond et al., 2007). These species use a stone hammer, often combined with a stone or wooden anvil, to crack open encased food sources such as nuts or shellfish (Barrett et al., 2018; Boesch & Boesch, 1990; Falótico & Ottoni, 2016; Luncz et al., 2017). This is widely considered to be a particularly sophisticated tool behavior due to its complexity and flexibility (Carvalho et al., 2008; Neadle et al., 2020; Visalberghi et al., 2017). It is also a very rare form of tool use across the primate order (Neadle et al., 2020), suggesting it may require specific conditions to emerge and be maintained. Primate stone tool use thus provides a valuable opportunity to investigate how advanced technological behaviors can arise and persist, serving as a model system for understanding early hominin stone technologies (Bandini et al., 2022).

It is widely accepted that, in addition to social and cognitive requisites, tool use likely requires certain environmental conditions to evolve (Koops & Sanz, 2022; Sanz & Morgan, 2013; Van Schaik et al., 1999). Evolutionary theories aimed at explaining the presence and absence of primate stone tool use have paid particular attention to the availability of local resources. Thus far, evidence regarding these theories has largely pertained to food resources, including food scarcity (Spagnoletti et al., 2012; Yamakoshi, 1998) and the availability or nutritional value of tool-extractable foods (Fox et al., 2004; Izar et al., 2022; Koops et al., 2013; Sanz & Morgan, 2013). Equivalent literature focused on raw tool materials is less developed. This is despite the knowledge that primate populations have access to and use different types of stones (e.g. across chimpanzees: Proffitt et al., 2022; across bearded capuchins: Visalberghi et al., 2007 vs. Falótico & Ottoni, 2016). Previous work investigating the frequency or spatial distribution of primate tool use sometimes discusses stone abundance or distribution of material (e.g. Almeida-Warren et al., 2022; Boesch et al., 1994; McGrew et al., 1997; Visalberghi et al., 2015). Only a small number of studies have considered the types of materials available. These have largely focused on how material properties may influence the accumulation and distribution of artifacts, with the aim of informing archaeological investigation (Reeves et al., 2023b, 2023a, 2024). However, raw material properties are not just important for shaping the archaeological record. Different materials may also provide tools of different quality, thereby influencing the foraging outcomes of tool use.

Tool quality can be defined in terms of performance – defined as how well a tool achieves an intended task – and durability – defined as the rate at which performance changes with use (Schunk, 2021). Differences in raw material performance may influence where tool use evolves. In theory, tool use should be more likely to evolve where it is highly profitable relative to alternative foraging behaviors (Rutz & St Clair, 2012; Sanz & Morgan, 2013). If local tool materials are highly efficient at extracting food, then this may decrease energetic costs and increase the relative profitability of tool use. Differences in material durability may also influence how tool use behavior shapes the technological landscape primates live in. The local accumulation of used tools has been suggested to promote social learning (Fragaszy et al., 2013a). Repeated small-scale transport of tools has also been shown to contribute to wide-scale distributions of tool materials away from their source, broadening the opportunities populations have for tool use (Reeves et al., 2021). Such processes may be hampered when local materials are not very durable, and do not last long under use (Reeves et al., 2023a).

Archaeological studies on hominin cutting-edge stone flakes suggest that both stone tool performance and durability can depend heavily on the raw material they are made of (Abrunhosa et al., 2019; Braun et al., 2009; Key et al., 2020). In comparison, the influence of raw material on the quality of primate percussive tools is poorly understood. While it is frequently asserted that primate tool efficacy or durability can be influenced by raw material, there is minimal quantifiable evidence of such effects (e.g., Sirianni et al., 2015; Visalberghi et al., 2007). Such assertions are commonly made in the context of tool selection behaviors, where functional differences between tools are either assumed as part of the experimental design, or inferred from observable tool preferences (Boesch & Boesch, 1983; Fragaszy et al., 2010; Visalberghi et al., 2009a, 2015). For instance, the fact that chimpanzees in the Taï National Park, Côte d’Ivoire, prefer stone hammers over wood when cracking hard nuts (Boesch & Boesch, 1983; Luncz et al., 2012; Sirianni et al., 2015), and the tendency for capuchins to transport harder versus softer stones to nut cracking sites (Visalberghi et al., 2007, 2009a, 2009b), may imply that these materials perform better in those contexts. However, preference alone cannot be used to definitively determine that materials differ in quality, as primates also exhibit cultural preferences which are not linked to tool performance (Luncz et al., 2012). Quantitative, experimental measures of tool quality are needed to further our understanding of this area.

Here, we provided experimental stone tool sets, varying in their raw material, to semi-wild bearded capuchins (*Sapajus libidinosus*) in Tietê Ecological Park, São Paulo, Brazil. These capuchins are frequently observed using stone tools to crack palm nuts (*Syagrus romanzoffiana*) (Luncz et al., 2024). Capuchins were provided with one of six tool sets at a time, each consisting of different combinations of anvil and hammer stone materials. Nut cracking behavior was recorded and scored for each tool set. We measured the performance and durability of tool materials using a number of metrics, including the probability of successfully cracking a nut, the hits required, and how long the tool lasts before it becomes unusable. Our results show how different raw materials used for percussive tools affect foraging performance in capuchin monkeys. This provides insight into how raw materials may affect the costs and benefits of tool-mediated foraging.

## METHODS

### Study Group

Data were obtained across ten days in April of 2017 from a population of 33 semi-wild bearded capuchins residing in the 14 km_2_ Tietê Ecological Park of São Paulo, Brazil. This population naturally uses stone tools to crack palm nuts (*Syagrus romanzoffiana*). We provided this group with experimental stone tools and naturally occurring oil palm nuts that we collected from their territory (Fig. 1).

**Figure 1:**
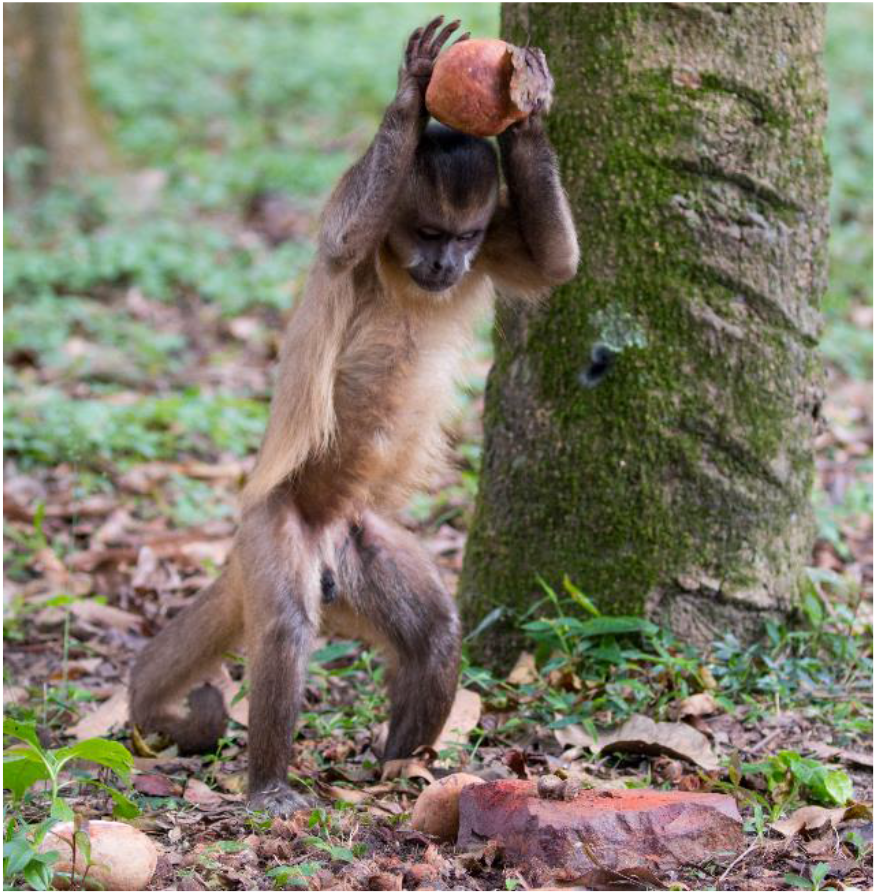
Capuchin nut cracking experimental set up. A semi-wild population of bearded capuchins (*Sapajus libidinosus*) was provided with one anvil and three hammer stones at a time (small, medium, and large), as well as locally collected palm nuts (*Syagrus romanzoffiana*). Photo credit: T.F.

### Stone Tool Raw Materials

The capuchins were provided with experimental tool sets differing in their combinations of raw materials (Table 1). Emphasis was placed on testing a wide variety of stone types, while limiting material combinations to those contextually relevant to hominin and primate tool activity. As such, materials were sourced globally from sites of nut cracking primates or early hominin tool use sites. Some anvils were quickly destroyed and replaced with different materials. These anvils are reported in the study to allow for discussion of differences in tool durability.

**Table 1.**
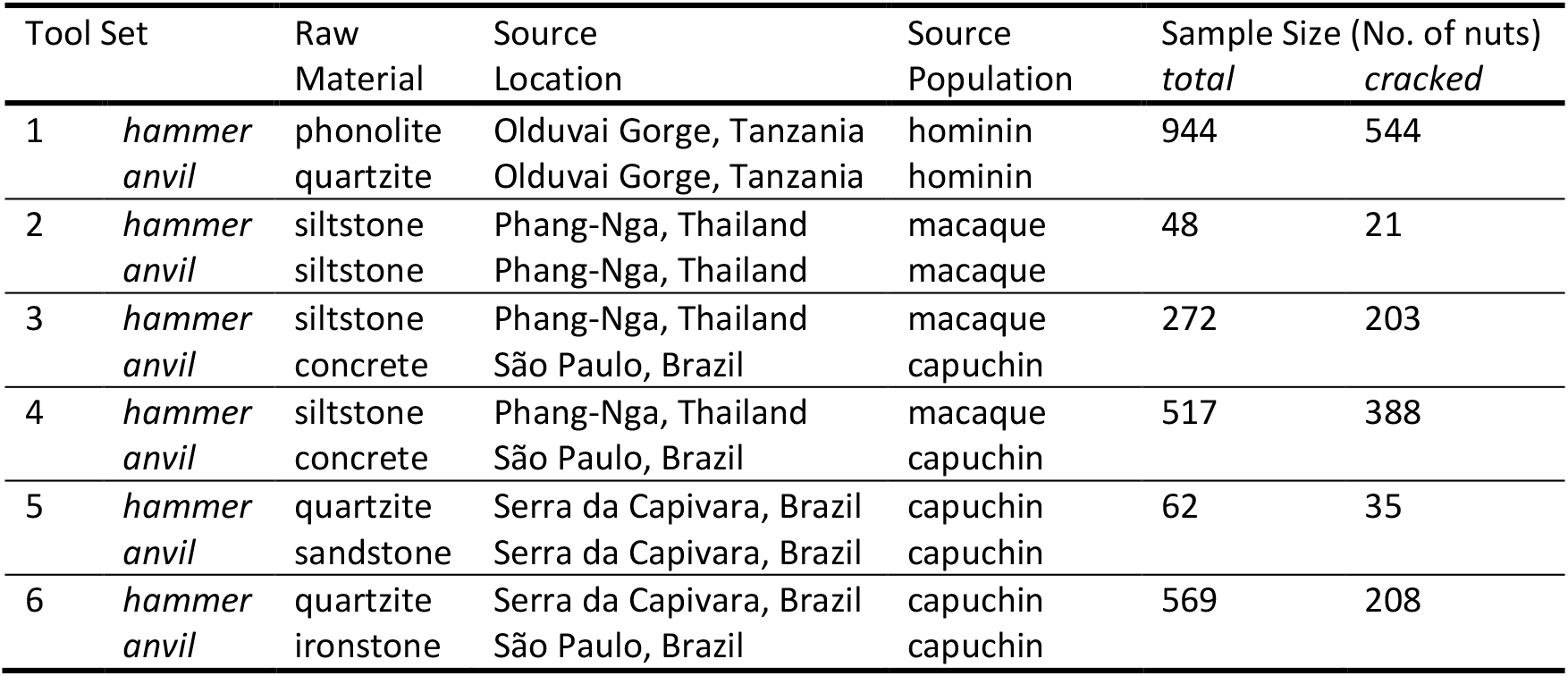
The raw material, source, and nut sample size for each tool set provided to capuchin monkeys. Sample sizes indicate the total number of nuts processed, and the number of these nuts which were cracked, after removing nuts processed by juveniles (n = 499 total) and nuts for which information was incomplete (n = 47).

One tool set was provided at a time, consisting of a stone anvil and three hammer stones. Each anvil was a flat tabular stone block approximately 16 to 30 cm across. All anvils were approximately 5 to 6 cm thick, except for the first concrete anvil (tool set 3), which was 2.5 cm thick (SI Table 1 for all tool sizes). The hammer stones in each tool set included a small (c. 300 g), medium (c. 400 g), and large (c.620 g) rounded cobble of identical material. This allowed each capuchin to choose their preferred size for nut cracking. The one exception was the ironstone-quartzite tool set (tool set 6), for which the large quartzite hammer stone fractured in half shortly into the study. As a result, the largest hammer stone in this tool set was approximately 400 g. A new tool set was provided when the material had degraded beyond the point of use, or when approximately 1000 nuts had been processed.

### Behavioral Observation

Nut cracking behaviour was recorded with a camcorder Canon Vixia HF R52 and a camera Canon EOS 70D. Recording took place from the moment a monkey approached the tool set to the moment the last individual left the tool set. From the videos, the following data was collected for each nut-cracking event: the tool set, weight of the hammerstone used by the capuchin, the number of hits applied to the nut, whether the nut was successfully cracked open, and the individual’s identity, age, and sex. This data was used to provide three measures of tool performance: the likelihood of successfully cracking a nut (cracking success), the number of hits required to crack a nut (hits-per-nut) and the number of hits required to obtain a food reward (hits-per-reward). Hits-per-nut excludes unsuccessful cracking attempts. Hits-per-reward is the combined number of hits in a successful cracking attempt, and the preceding unsuccessful attempts, providing a measure of overall efficiency. A measure of accumulating ‘use’ was also generated: the cumulative number of hammer strikes applied to the anvil prior to the given nut, including hammer strikes performed by juveniles. This provides an assessment of how much a tool set had been used at each point in the study, thereby enabling assessment of how tool performance changes over time.

### Statistical Modelling

Differences in tool performance and durability across materials was assessed using generalized liner mixed models (GLMMs). Models were generated using the package ‘glmmTMB’ (Brooks et al., 2017) in R (v 4.3.1; R Core Team, 2018). Three models were used, each addressing a different performance metric: 1. cracking success (yes/no), 2. hits-per-nut, and 3. hits-per-reward were compared across tool sets. The sample size for cracking success (model 1) corresponds to the total number of nuts processed (n = 2302), and includes 12 individuals (8 males, 4 females). The sample size for hits-per-nut (model 2) and hits-per-food reward (model 3) corresponds to the number of nuts successfully cracked (n = 1343), and includes 11 individuals (8 males, 3 females). Nuts processed by juveniles (n = 499 nuts, 8 male juveniles < 5 years old) were excluded from analysis, as these individuals may still be developing their nut cracking abilities. The tool sets with the siltstone and sandstone anvils (tool sets 2 and 5 respectively) were also excluded due to low sample sizes due to the rapid destruction of their anvils (Table 1 for sample size of each material).

All models had the same predictor variables: material, use, hammer weight, and sex. There were two variables of interest: ‘material’ as a categorical variable reflecting the tool set, and ‘use’ as measured in cumulative prior hammer strikes. An interaction between material and use was included to test if the rate at which tool performance changes with use depends on the material the tool is made of. Hammer weight and sex were included as control variables. Also included was a random effect of subject ID on the intercept, and on the slope of the material effect. Inclusion of such a random slope is shown to reduce type I errors providing a conservative estimate of significance (Matuschek et al., 2017). When assessing hits-per-reward (model 3), hammer weight reflects the mean weight of hammers used across the successful nut and its unsuccessful predecessors. Hammer weight and use were scaled by subtracting from the mean and dividing by the standard deviation. The cracking success model (model 1) used a binomial distribution and logit link function. The hits-per-nut and hits-per-reward models (models 2 and 3) used a zero-truncated negative binomial distribution to correct for overdispersion and a right skew, with a log link function.

All models performed well when tested for homoscedasticity of residuals, overdispersion, collinearity of predictor variables, and model stability (SI ‘Model Diagnostics’). Homoscedasticity was assessed by visually inspecting plots of residual values against predicted values. Residuals were found to vary equally across the range of predicted values. Dispersion was assessed using the ‘DHARMa’ package (Hartig, 2022). The dispersion ratios were close to 1 (0.89 to 1.02 depending on the model) and models were not significantly over or underdispersed (*P* > 0.05). Collinearity was low with VIF values of 1.66 or less. Model stability was assessed by dropping one individual at a time, with all variables of interest found to be reasonably stable. The binomial cracking success model (model 1) was also checked for complete separation by visually checking sample sizes across groups.

The significance of each model was tested by comparing it to a null model using the Anova function in R. Thereafter, the significance of the effect of each individual predictor variable was assessed by dropping the variable from the model, and comparing the resulting reduced model to the full model. This reduced versus full model comparison was done, one at a time, for material, use, the interaction of material and use, and hammer weight. The predicted influence of material and use was plotted for each response variable using the predict function.

### Ethical Note

This research complied with protocols approved by the Animal Research Ethical Committee of the Institute of Psychology, University of São Paulo (CEUA 3036140715), fully adhered to Brazilian law under authorization from agencies IBAMA/ICMBio 37609-6 and CNPq 001375/2015-60, complied with the American Society of Primatologists Principles for the Ethical Treatment of Non-Human Primates, and adheres to the ARRIVE guidelines. No stress or adverse effects were observed or anticipated from the procedures performed in this study. All subjects were free-living animals participating at will with the experiment. This population was habituated to and comfortable around human presence. The nut species and size of tools provided reflected those naturally available to the species, and the observed nut cracking was within their typical behavioral repertoire.

## RESULTS

### Material Durability

Materials differed in how easily they fractured during use. For hammerstones, only one large quartzite hammer fractured, splitting in half after 166 strikes. Anvils fractured much more readily. The siltstone and sandstone anvils quickly fractured into a large number of small pieces (> 30 each) in less than 84 and 129 strikes, respectively. They were thus removed from the study. The first concrete anvil had its first large piece break off after only 16 strikes, and after 700 strikes it had been broken into six pieces (7.5-14 cm long). The ironstone anvil developed a large depression and eventually split in half after 536 strikes. Both halves remained viable anvils. One half was retained for the remaining duration of the study, accumulating a further 974 hammer strikes (1510 total for the anvil). The quartzite and second concrete anvils were the only anvils that remained largely intact by the end of the study, after sustaining 1758 and 1052 hammer strikes, respectively. Although both anvils lost pieces along their outer edge up to 14 cm in length, these pieces were generally thin and did not alter the overall size of the anvils by more than a few centimeters in either direction (SI Fig. 2).

### Model Overview

GLMM analysis reveals that tool performance is influenced by the tool attributes included in the models. All three models differed significantly from their respective null model (SI Table 2 for all model comparisons). Further full versus reduced model comparison for all three models did not find support for the presence of an interaction between material and use. This indicates the rate at which tool performance changes with use does not differ depending on the material the tool is made of. However, material, hammer stone weight, and the prior use of the tool all had important effects on the aspects of tool performance investigated. These are discussed in detail below. We report model outputs (effect sizes and predicted values) using the reduced version of each of the three performance models, where there is no interaction (full model results in SI Table 3). Unless stated otherwise, all predicted values assume a male capuchin, an unused tool (prior uses = 0), and a hammer weight of 518 g (the mean for this study).

### Cracking Success

Overall, only slightly more than half of all nuts were cracked successfully (58%). Unsuccessful nut cracking events were largely the result of the nut flying off the anvil when struck. The likelihood of cracking a given nut depended heavily on the tool material and hammer weight. Material was shown to have a highly significant effect (model 1: full vs. reduced model comparison: *X*_*2*6_ = 25.9, *P* < 0.001), with nuts being cracked more consistently using some materials over others (effect size E between -1.06 and 1.04; Table 2). The best performing material was tool set 4 (i.e. concrete2-siltstone: the second concrete anvil paired with siltstone hammerstones). This material was over three times more likely to crack a given nut than the worst material (tool set 6: ironstone-quartzite) (predicted likelihoods of 0.83 versus 0.25; Fig. 2a). Larger hammers were also significantly more effective at cracking nuts (*X*_*2*1_ = 5.5, *P* < 0.05). The likelihood of successfully cracking a nut increased significantly with hammer weight (E ± SD = 0.11 ± 0.051). When comparing the smallest hammer (c. 260 g) to the largest (c. 650 g), the chance of cracking a nut rose by 20% (predicted likelihood of 0.52 vs 0.62; SI Fig. 7). While 20% is still a notable difference, this is much less than the 300% difference between materials. The influence of hammer weight may be somewhat muted by the fact that capuchins were able to choose their preferred hammer. Capuchins chose the largest available hammer 77% of the time. Tools did not get better or worse at cracking nuts over time: the full model did not differ from the reduced model where prior use was removed (*X*_*2*4_ = 6.4, *P* = 0.17).

**Figure 2:**
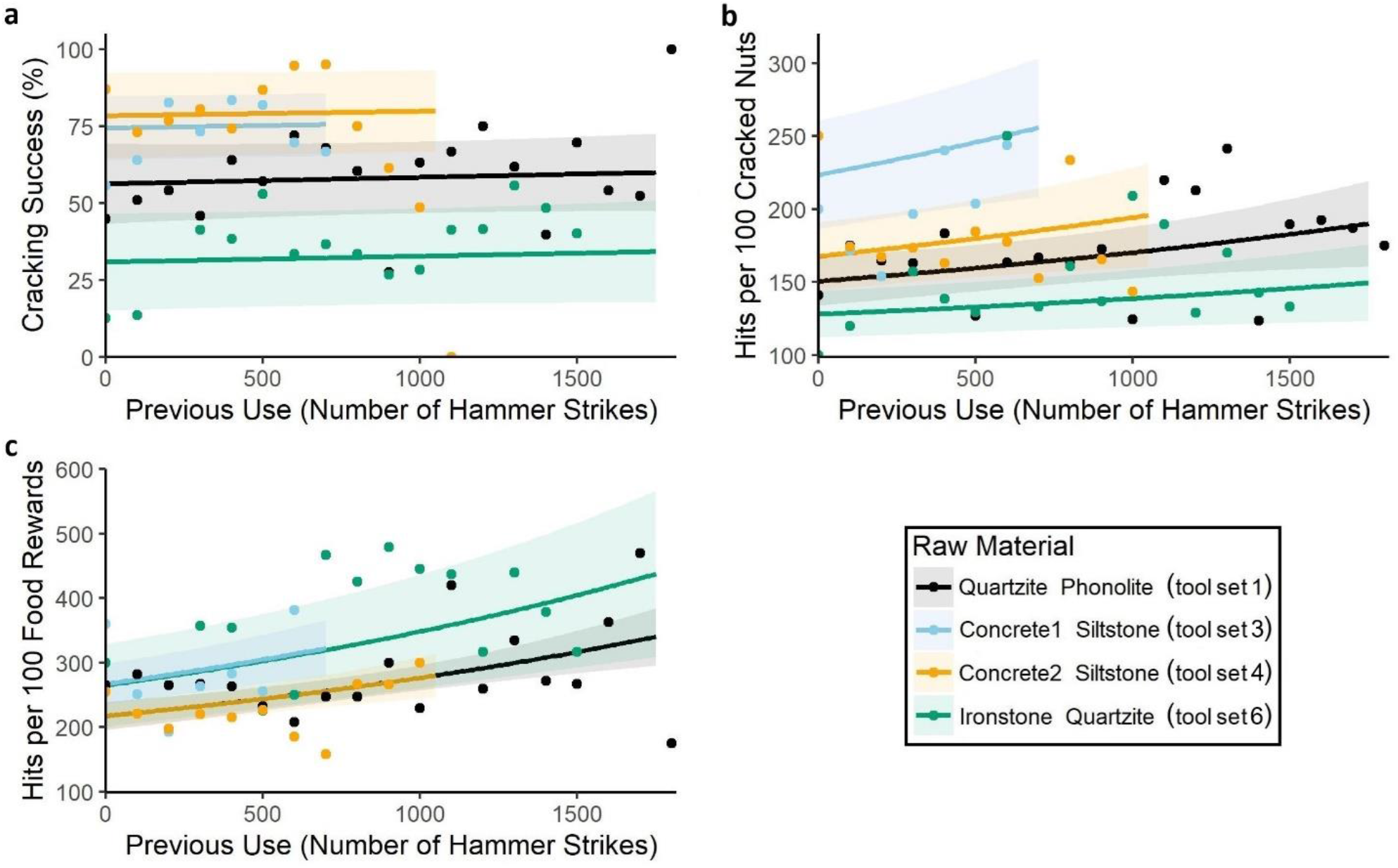
Influence of raw material and prior use on tool performance. Tool performance as measured by the likelihood of successfully cracking a nut (a, model 1), hits to open a nut (b, model 2), and hits to obtain a food reward (c, model 3). Performance varies across tool sets of different raw material (listed as anvil then hammer stone) and with increasing use. Each point indicates the observed data, averaged across 100 hammer strikes. Three outlaying points are excluded to improve interpretation of the slope (visible in SI Fig. 6). Lines reflect model predictions, assuming a male capuchin and hammer weight of 518 g, with no interaction between raw material and use. Shaded areas reflect one standard error either side of the estimate.

**Table 2.** GLMM outputs regarding cracking success, hits-per-nut and hits-per-reward (interaction between material and use removed). Estimates for hammer weight and use have been scaled to reflect change per 100 grams and 100 cumulative hammer strikes respectively. The base material was tool set 1, which had a quartzite anvil and phonolite hammerstones (quartzite-phonolite). Material effects are from tool set 3 (concrete1-siltstone), tool set 4 (concrete2-siltstone) and tool set 6 (ironstone-quartzite).

| Parameter | Model 1: Cracking-Success |  |  | Model 2: Hits-Per-Nut |  |  | Model 3: Hits-Per-Reward |  |  |
| --- | --- | --- | --- | --- | --- | --- | --- | --- | --- |
|  | Estimate | (SD) | Z | Estimate | (SD) | Z | Estimate | (SD) | Z |
| Intercept | 0.31 | (0.24) | 1.26 | -0.25 | (0.15) | -1.70 | 0.44 | (0.077) | 5.67 |
| Material |  |  |  |  |  |  |  |  |  |
| *Tool Set 3* | 0.81 | (0.21) | 3.87 | 0.86 | (0.14) | 6.02 | 0.36 | (0.12) | 2.96 |
| *Tool Set 4* | 1.04 | (0.37) | 2.84 | 0.28 | (0.16) | 1.81 | -0.00035 | (0.10) | -0.0034 |
| *Tool Set 6* | -1.06 | (0.30) | -3.54 | -0.58 | (0.24) | -2.41 | 0.34 | (0.19) | 1.80 |
| Use | 0.0087 | (0.013) | 0.67 | 0.032 | (0.012) | 2.66 | 0.041 | (0.0088) | 4.65 |
| Hammer Weight | 0.11 | (0.051) | 2.07 | -0.27 | (0.039) | -6.78 | -0.24 | (0.037) | -6.58 |
| Sex Female | -1.49 | (0.51) | -2.90 | 0.67 | (0.31) | 2.18 | 0.87 | (0.22) | 4.05 |

### Hits-Per-Nut

Tool material, hammer weight, and prior use were all important variables in determining the number of hits required to crack a nut. Tools made of different materials varied considerably in the number of hits required to open a nut (model 2: E between -0.58 and 0.86; full vs. reduced model comparison: *X*_*2*6_ = 25.5, *P* < 0.001). Larger hammers also required fewer hits than smaller hammers (E ± SD = -0.27 ± 0.039; *X*_*2*1_ = 39.8, *P* < 0.001). While the majority of nuts were opened in one (59%) or two (24%) hits, capuchins in our study cracked as many as 99 nuts in one sitting. In these circumstances, small differences in effort on a per-nut basis may have large cumulative effects. We thus report our results in terms of hits per 100 nuts, as calculated by multiplying the predicted hits per nut by 100. Comparing the predicted number of hits across different materials and hammer weights shows both these factors are similarly important. The worst performing material (tool set 3: concrete1-siltstone) required almost twice as many hits compared to the best material (tool set 6: ironstone-quartzite) (predicted hits per 100 nuts: 223 vs. 128; Fig. 2b). Similarly, the smallest hammer required slightly less than twice the number of hits compared to the largest hammer (predicted hits: 240 vs. 147). Tools also appeared to perform worse as they accumulated use. The predicted number of hits increased from 167 to 220 when comparing an unused tool to a heavily used tool (0 vs. 1750 prior strikes; E ± SD = 0.032 ± 0.012; *X*_*2*4_ = 12.3, *P* < 0.05). However, this result appears to be driven by one tool set. The comparatively thin concrete anvil in tool set 3 split in half partway through the study. When nuts were placed along the fracture, the anvil would move when struck, likely dissipating the force of the hammerstone. Rerunning the full versus reduced model comparison without tool set 3 showed performance did not significantly degrade with use (*X*_*2*3_ = 4.7, *P* = 0.19).

### Hits-Per-Reward

The energetic efficiency of nut cracking varied according to tool material, hammer weight, and prior use. Full-versus reduced model comparisons show these effects were significant. As with the hits-per-nut model, results for this model are discussed in terms of hits per 100 food rewards. Materials differed in their efficiency, although this difference was not as strong as for the other performance metrics (model 3: E between 0 and 0.36; *X*_*2*6_ = 17.6, *P* < 0.01). Efficiency increased by 18% from the worst materials (tool sets 3 and 6: concrete1-siltstone and ironstone-quartzite; c. 265 predicted hits) to the best materials (tool sets 1 and 4: quartzite-phonolite and concrete2-siltstone; 217 hits each; Fig. 2c). Hammer weight was important for efficiency, with larger hammers requiring substantially fewer hits (E ± SD = -0.24 ± 0.037; *X*_*2*1_ = 40.7, *P* < 0.001). Efficiency increased by 45% from the smallest (366 hits) to the largest (203 hits) hammer. Tools became less efficient over time as they accumulated use (E ± SD = 0.041 ± 0.0088; *X*_*2*4_ = 24.6, *P* < 0.001). The predicted number of hits per 100 food rewards increased by 61% (242 vs. 390 hits) when comparing an unused tool (0 prior strikes) to a heavily used tool (1750 prior strikes). Unlike in the hits-per-nut model, the effect of use in this model does not appear to be due solely to tool set 3 (concrete1-siltstone). Even if this material is removed, there is a significant difference between the full model and a reduced model which excludes an effect of use (X_23_ = 20.6, *P* < 0.001).

## DISCUSSION

Most studies investigating the environmental context of stone tool use in non-human primates have focused heavily on the quality and availability of extractable food resources (Fox et al., 2004; Izar et al., 2022; Koops et al., 2013; Sanz & Morgan, 2013; Spagnoletti et al., 2012; Yamakoshi, 1998). There has been comparatively little emphasis on the effect of available raw tool material on foraging efficiency. Here we investigated if raw material affects the performance and durability of tools used by non-human primates for nut cracking.

All raw materials provided to the monkeys were successfully used to crack nuts. However, raw materials differed significantly in performance. The strongest difference was in the likelihood of successfully cracking open a nut. The best performing material was a concrete anvil paired with siltstone hammerstones (tool sets 3 and 4). With this material capuchins were able to crack twice as many of the nuts provided compared to the worst performing material (tool set 6: ironstone anvil with quartzite hammerstones). Failure to crack a nut was typically due to the nut flying off the anvil when struck, suggesting this is an issue of nut stability. In this study, the nuts provided were round, making stability on the anvil an issue. However, other natural resources may not be so sensitive to stability issues. In a competitive species where food scrounging is common (Coelho et al., 2015; Vogel, 2005), the nut may not be retrievable once lost, which might result in a lower overall food yield. The importance of nut stability for cracking efficiency is widely recognized in the literature, but only in the context of anvil pitting (Fragaszy et al., 2010, 2013b; Haslam et al., 2014; Liu et al., 2011). Anvils in this study accumulated surface damage, including shallow, wide depressions that could be described as pits (Luncz et al., 2022 for details). However, nut stability did not change over time, indicating raw materials can differ in their nut stability independent of their capacity to form pits. Anvils with higher surface friction may provide higher nut stability. In this study the concrete anvils had a rough surface texture and also was shown to exhibit the greatest nut stability. In comparison, the ironstone anvil had the lowest nut stability and a surface which damaged easily. This surface may give way partially when struck, reducing the friction on the nut. Determining which stone properties drive differences in tool performance is a complex task, and a matter of ongoing investigation (e.g. for hominin tools; Lerner et al., 2007; Lin et al., 2023; Luncz et al., 2022).

Raw materials also differed significantly in the number of hits required to crack open a nut. This performance metric is often used to infer energetic efficiency of nut cracking (Boesch et al., 2017; Falótico et al., 2024; Luncz et al., 2024). However, in this study, the number of hits per nut appeared to be influenced by nut stability. The hits per nut for a given material was inversely proportional to its cracking success, the latter of which was directly influenced by nut stability as discussed above. When nut stability is low, then there is a high chance of losing a nut each time it is struck. Harder nuts which require more hits are therefore more likely to be lost, and will not contribute to the hits per nut for that material. This can make low-stability materials appear more ‘efficient’ than they are. Instead, efficiency can be accurately described by the number of hits required to obtain a food reward. This performance metric differed significantly across materials, with an approximately 20% difference between the best materials (tool set 1: quartzite anvil and phonolite hammerstones, and tool set 4: thicker concrete anvil and siltstone hammerstones) and the worst materials (tool sets 3: thinner concrete anvil and siltstone hammerstones, and tool set 6: ironstone anvil and quartzite hammerstones).

Comparison between tool sets suggests that multiple factors influence efficiency. Tool sets 1 and 4 had similarly high efficiencies. Tool set 1 achieved its high efficiency by having a high nut stability, as measured by likelihood of cracking a nut. In comparison, tool set 4 achieved a similar efficiency by cracking nuts in a small number of hits, suggesting this material had a highly effective hammer strike. Quartzite (the anvil for tool set 4) tends to be harder than concrete (anvil in tool set 1) and may absorb less force (respective Leeb hardness of 445-729 vs. 180-460 HL; Alberti et al., 2013; Kovler et al., 2018). Stone hardness may thus be an important predictor for hammer strike effectiveness. Efficiency is also influenced by tool mass. Increases in hammer mass significantly reduced the number of hits required to open a nut. Comparison between tool sets 3 and 4 also reveal an influence of anvil mass: both tool sets had a concrete anvil with siltstone hammers, but the anvil in tool set 4 was twice as thick and required far fewer hits per nut. While prior research has identified an influence of hammer mass on nut cracking (chimpanzees; Boesch et al., 2017), anvil mass has largely been disregarded.

The patterns of raw material performance observed here have implications for the conditions under which tool use may be expected to evolve. All raw materials provided to the monkeys were successfully used to crack nuts, suggesting a wide array of materials are suitable for this tool use behavior. This aligns with prior studies, where percussive artifacts used by extant primates can consist of a wide range of stone types (Reeves et al., 2023b, 2024). It is also consistent with the fact that stone-mediated nut cracking has evolved in a wide variety of material landscapes (e.g. capuchins; Visalberghi et al., 2007; macaques; Proffitt et al., 2018; chimpanzees; Proffitt et al., 2022). As such, the type of stones available to a primate population may not present a major barrier to the emergence of tool use, and is unlikely on its own to explain the absence or presence of tool use. However, differences in raw material performance may subtly augment the likelihood of percussive behaviors emerging in a given population. The relatively profitability hypothesis states that tool use should only be retained where it is more profitable than foraging without tools (Rutz & St Clair, 2012; Sanz & Morgan, 2013). Raw materials in this study differed in the food yield and energetic efficiency they confer: both factors are likely to affect the profitability (i.e. cost to benefit ratio) of tool use. Furthermore, stone tool use is predominantly observed on the ground in capuchins (Falótico & Ottoni, 2023), macaques (Luncz et al., 2017; Malaivijitnond et al., 2007), and chimpanzees (Boesch & Boesch, 1983). Many primate populations face ground-based predation (Ferrari, 2009; Zuberbühler & Jenny, 2002), which has been theorized to limit opportunities for terrestrial tool use (Monteza-Moreno et al., 2020). Where ground-based predation is prevalent, the trade-off between tool use and predation risk may be partially alleviated by having access to highly efficient raw materials. Similar evolutionary pressures have been suggested for hominins, which faced predation by large mammals and may have reduced this predation risk by using tool materials which allowed for more efficient foraging techniques (Caruana, 2020).

In addition to tool performance, our results revealed notable differences in tool durability depending on raw material. Sedimentary anvils saw the highest rates of fracturing. The sandstone and siltstone anvils in particular were rapidly destroyed, highlighting the brittle nature of these stones. The anvils with the least damage were the quartzite and large concrete anvil: materials respectively known for their hardness (Howard, 2005) and valued as durable construction material. This pattern of durability matches what we see across wild primate populations. Natural sandstone anvils used by capuchins can accrue large pits and lose large proportions of their mass in short periods of time (Haslam et al., 2014). In comparison, quartzite anvils used by captive chimpanzees show limited fracturing (Arroyo et al., 2016).

Differences in material durability have implications for the processes which shape technological landscapes. The accumulation of used tools may facilitate learning in naïve individuals by encouraging them to interact with tools that have visible signs of use (Fragaszy et al., 2013a). Furthermore, through repeated short-scale transport of stones to tool use sites, populations may expand the reach of tool material across their home range (Reeves et al., 2021). Through processes such as these, primates may unintentionally improve opportunities for future generations to access and learn how to use tools. However, such processes may be diminished where tools are less durable, with limited potential for re-use (Reeves et al., 2023a). The results from this study broadly suggest that sedimentary stones may not last as long. In landscapes dominated by sedimentary material, tools may not be transported as far from their source material before breaking, and naïve individuals may have less frequent access to used tools that present salient social cues. However, it should be noted that the influence of raw material durability on the technological landscape is likely augmented by other factors, such as tool size and abundance. In Fazenda Boa Vista, large sedimentary anvils used by capuchins can last for several years despite rapidly losing mass if used frequently (Haslam et al., 2014). This longevity can be attributed to their large starting mass, and the fact that nut cracking activity is spread across numerous anvils, reducing the intensity of use on a single tool. In these conditions, the low durability of the material may facilitate social learning, instead of impeding it, because these anvils rapidly acquire visible damage.

This study provides evidence that raw materials differ in their performance and durability as primate nut cracking tools. However, these results may depend on the primate species in question. Recent work comparing tool behavior across capuchins, macaques, and chimpanzees shows that the two-handed nut cracking technique used by capuchins is relatively imprecise (Luncz et al., 2024). The hammer stone frequently impacts the anvil causing damage to the tools, and the nut frequently flies off the anvil when struck. In comparison, macaques (and to a lesser extent chimpanzees) have a high precision one-handed striking technique which produces significantly less tool damage, and allows them to use their free hand to shield and catch loose nuts. Based on this comparison, the nut stability and resistance to fracturing afforded by different raw materials may be more important for capuchins than for more precise nut-cracking species.

Overall, this study demonstrates that different raw materials do not produce equivalent tools. Tool performance differs due to the nut stability afforded by the material, and durability depends on how easily materials fracture. Differences across materials may also help shape how tool use evolves after its emergence, including by altering the dynamic processes shaping technological landscapes. These results affirm that raw material properties should be a central consideration when investigating the origins of tool use, not just when assessing lithic evidence, but also for behavioral and selective processes in living species.

## ACKNOWLEDGEMENTS

We thank the Tietê Ecological Park for enabling our research. We thank Henrique P. Rufo for the identification of the monkeys, and Tatiane Valença for the coding of the videos. Raw material collection at Olduvai was authorized by the Commission for Science and Technology (COSTECH), the Ngorongoro Conservation Area Authority, and the Department of Antiquities, Tanzania under the following COSTECH permits (ORACEAF, ERC-StG 283366). This research was funded by the German Primate Center and the Max Planck Institute for Evolutionary Anthropology. T.F. was supported by the São Paulo Research Foundation (2013/05219-0 and 2018/01292-9). I.T. is supported by an ERC-Advanced Grant (BICAEHFID, No. 832980). T.P. was supported by a British Academy Fellowship (Project Number: 542133) and a Leakey Foundation Grant and is currently supported by grant CEECINST/00052/2021 funded by the Portuguese Foundation for Science and Technology, Portugal.

## AUTHOR CONTRIBUTIONS

L.V.L and T.P conceived the study. L.V.L and T.F collected primate data. I.T. provided raw material from Olduvai Gorge. T.D.R.O. analyzed the data and produced the figures with contributions from L.V.L., J.R. and N.E.S. The paper and Supplementary information were written by T.D.R.O and L.V.L. with contributions from J.R., T.P., N.E.S., T.F. and I.T.

## REFERENCES

Abrunhosa, A., Pereira, T., Márquez, B., Baquedano, E., Arsuaga, J. L., & Pérez-González, A. (2019). Understanding Neanderthal technological adaptation at Navalmaíllo Rock Shelter (Spain) by measuring lithic raw materials performance variability. Archaeological and Anthropological Sciences, 11, 5949–5962. 10.1007/s12520-019-00826-3

Alberti, A. P., Gomes, A., Trenhaile, A., Oliveira, M., & Horacio, J. (2013). Correlating river terrace remnants using an Equotip hardness tester: An example from the Miño River, northwestern Iberian Peninsula. Geomorphology, 191, 59–70. 10.1016/j.geomorph.2013.03.017

Almeida-Warren, K., Camara, H. D., Matsuzawa, T., & Carvalho, S. (2022). Landscaping the behavioural ecology of primate stone tool use. International Journal of Primatology, 43, 885–912. 10.1007/s10764-022-00305-y

Arroyo, A., Hirata, S., Matsuzawa, T., & De La Torre, I. (2016). Nut cracking tools used by captive chimpanzees (Pan troglodytes) and their comparison with early stone age percussive artefacts from Olduvai Gorge. PLoS ONE, 11(11). 10.1371/journal.pone.0166788

Bandini, E., Harrison, R. A., & Motes-Rodrigo, A. (2022). Examining the suitability of extant primates as models of hominin stone tool culture. Humanities and Social Sciences Communications, 9, 74. 10.1057/s41599-022-01091-x

Barrett, B. J., Monteza-Moreno, C. M., Dogandžić, T., Zwyns, N., Ibáñez, A., & Crofoot, M. C. (2018). Habitual stone-tool-aided extractive foraging in white-faced capuchins, Cebus capucinus. Royal Society Open Science, 5, 181002. 10.1098/rsos.181002

Boesch, C., & Boesch, H. (1983). Optimisation of nut-cracking with natural hammers by wild chimpanzees. Behaviour, 83, 265–268.

Boesch, C., & Boesch, H. (1990). Tool use and tool making in wild chimpanzees. Folia Primatologica, 54(1–2), 86–99. 10.1159/000156428

Boesch, C., Bombjaková, D., Boyette, A., & Meier, A. (2017). Technical intelligence and culture: Nut cracking in humans and chimpanzees. American Journal of Physical Anthropology, 163(2), 339–355. 10.1002/ajpa.23211

Boesch, C., Marchesi, P., Marchesi, N., Fruth, B., & Joulian, F. (1994). Is nut cracking in wild chimpanzees a cultural behaviour? Journal of Human Evolution, 26(4), 325–338. 10.1006/jhev.1994.1020

Braun, D. R., Plummer, T., Ferraro, J. V., Ditchfield, P., & Bishop, L. C. (2009). Raw material quality and Oldowan hominin toolstone preferences: evidence from Kanjera South, Kenya. Journal of Archaeological Science, 36(7), 1605–1614. 10.1016/j.jas.2009.03.025

Brooks, M. E., Kristensen, K., Benthem K. J. van, Magnusson, A., Berg, C. W., Nielsen, A., Skaug, H. J., Maechler, M., & Bolker, B. M. (2017). glmmTMB Balances Speed and Flexibility Among Packages for Zero-inflated Generalized Linear Mixed Modeling. The R Journal, 9(2), 378–400. 10.32614/RJ-2017-066

Canale, G. R., Guidorizzi, C. E., Kierulff, M. C. M., & Gatto, C. A. F. R. (2009). First record of tool use by wild populations of the yellow-breasted capuchin monkey (Cebus xanthosternos) and new records for the bearded capuchin (Cebus libidinosus). Americal Journal of Primatology, 71, 366–372. 10.1002/ajp.20648

Caruana, M. V. (2020). Exploring the influence of predation risks on Oldowan tool use in South Africa. Journal of Field Archaeology, 45(8), 608–620. 10.1080/00934690.2020.1813394

Carvalho, S., Cunha, E., Sousa, C., & Matsuzawa, T. (2008). Chaînes opératoires and resource-exploitation strategies in chimpanzee (Pan troglodytes) nut cracking. Journal of Human Evolution, 55(1), 148–163. 10.1016/j.jhevol.2008.02.005

Coelho, C. G., Falótico, T., Izar, P., Mannu, M., Resende, B. D., Siqueira, J. O., & Ottoni, E. B. (2015). Social learning strategies for nut-cracking by tufted capuchin monkeys (Sapajus spp.). Animal Cognition, 18, 911–919. 10.1007/s10071-015-0861-5

De Beaune, S. A. (2004). The invention of technology: Prehistory and cognition. Current Anthropology, 45(2), 139–162. 10.1086/381045

Falótico, T., Macedo, A. C., de Jesus, M. A., Espinola, T., & Valença, T. (2024). Nut-cracking success and efficiency in two wild capuchin monkey populations. Royal Society Open Science, 11(6). 10.1098/rsos.240161

Falótico, T., & Ottoni, E. B. (2016). The manifold use of pounding stone tools by wild capuchin monkeys of Serra da Capivara National Park, Brazil. Behaviour, 153(4), 421–442. 10.1163/1568539X-00003357

Falótico, T., & Ottoni, E. B. (2023). Greater tool use diversity is associated with increased terrestriality in wild capuchin monkeys. American Journal of Biological Anthropology, 181(2), 312–317. 10.1002/ajpa.24740

Ferrari, S. F. (2009). Predation risk and antipredator strategies. In K B Garber, P.A., Estrada, A., Bicca-Marques, J.C., Heymann, E.W. Strier (Ed.), South American Primates. Springer. 10.1007/978-0-387-78705-3_10

Fox, E. A., Van Schaik, C. P., Sitompul, A., & Wright, D. N. (2004). Intra-and interpopulational differences in orangutan (Pongo pygmaeus) activity and diet: Implications for the invention of tool use. American Journal of Physical Anthropology, 125(2), 162–174. 10.1002/ajpa.10386

Fragaszy, D. M., Biro, D., Eshchar, Y., Humle, T., Izar, P., Resende, B., & Visalberghi, E. (2013a). The fourth dimension of tool use: Temporally enduring artefacts aid primates learning to use tools. Philosophical Transactions of the Royal Society B: Biological Sciences, 368, 20120410. 10.1098/rstb.2012.0410

Fragaszy, D. M., Greenberg, R., Visalberghi, E., Ottoni, E. B., Izar, P., & Liu, Q. (2010). How wild bearded capuchin monkeys select stones and nuts to minimize the number of strikes per nut cracked. Animal Behaviour, 80(2), 205–214. 10.1016/j.anbehav.2010.04.018

Fragaszy, D. M., Liu, Q., Wright, B. W., Allen, A., Brown, C. W., & Visalberghi, E. (2013b). Wild bearded capuchin monkeys (Sapajus libidinosus) strategically place nuts in a stable position during nut-cracking. PLoS ONE, 8(2), e56182. 10.1371/journal.pone.0056182

Hartig, F. (2022). DHARMa: residual diagnostics for hierarchical (multi-level/mixed) regression models. https://cran.r-project.org/web/packages/DHARMa/vignettes/DHARMa.html

Haslam, M., Cardoso, R. M., Visalberghi, E., & Fragaszy, D. M. (2014). Stone anvil damage by wild bearded capuchins (Sapajus libidinosus) during pounding tool use: A field experiment. PLoS ONE, 9(11), e111273. 10.1371/journal.pone.0111273

Howard, J. L. (2005). The quartzite problem revisited. The Journal of Geology, 113(6), 707–713. 10.1086/449328

Izar, P., Peternelli-dos-Santos, L., Rothman, J. M., Raubenheimer, D., Presotto, A., Gort, G., Visalberghi, E. M., & Fragaszy, D. M. (2022). Stone tools improve diet quality in wild monkeys. Current Biology, 32(18), 4088–4092.e3. 10.1016/j.cub.2022.07.056

Key, A., Proffitt, T., & de la Torre, I. (2020). Raw material optimization and stone tool engineering in the Early Stone Age of Olduvai Gorge (Tanzania). Journal of the Royal Society Interface, 17, 20190377. 10.1098/rsif.2019.0377

Koops, K., McGrew, W. C., & Matsuzawa, T. (2013). Ecology of culture: Do environmental factors influence foraging tool use in wild chimpanzees, Pan troglodytes verus? Animal Behaviour, 85(1), 175–185. 10.1016/j.anbehav.2012.10.022

Koops, K., & Sanz, C. (2022). Progress and prospects in primate tool use and cognition. In B. L. Schwartz & M. J. Beran (Eds.), Primate Cognitive Studies (pp. 238–259). Cambridge University Press. 10.1017/9781108955836.010

Kovler, K., Wang, F., & Muravin, B. (2018). Testing of concrete by rebound method: Leeb versus Schmidt hammers. Materials and Structures, 51, 138. 10.1617/s11527-018-1265-1

Langley, M. C., & Suddendorf, T. (2022). Archaeological evidence for thinking about possibilities in hominin evolution. Philosophical Transactions of the Royal Society B: Biological Sciences, 377(1866). 10.1098/rstb.2021.0350

Lerner, H., Du, X., Costopoulos, A., & Ostoja-Starzewski, M. (2007). Lithic raw material physical properties and use-wear accrual. Journal of Archaeological Science, 34(5), 711–722. 10.1016/j.jas.2006.07.009

Lima, G. C. B., Lacerda, J. C., Taynor, R., Araújo, M., Bezerra, B. M., & Souza-Alves, J. P. (2024). A new addition to the toolbox: stone tool use in blonde capuchin monkeys (Sapajus flavius). Primates, 65(5), 383–389. 10.1007/s10329-024-01143-7

Lin, S. C., White, L. T., Jatmiko Julianto, I. M. A., Tocheri, M. W., & Sutikna, T. (2023). Characterising the stone artefact raw materials at Liang Bua, Indonesia. Journal of Paleolithic Archaeology, 6, 22. 10.1007/s41982-022-00133-9

Liu, Q., Fragaszy, D. M., Wright, B., Wright, K., Izar, P., & Visalberghi, E. (2011). Wild bearded capuchin monkeys (Cebus libidinosus) place nuts in anvils selectively. Animal Behaviour, 81(1), 297–305. 10.1016/j.anbehav.2010.10.021

Luncz, L. V., Arroyo, A., Falótico, T., Quinn, P., & Proffitt, T. (2022). A primate model for the origin of flake technology. Journal of Human Evolution, 171, 103250. 10.1016/j.jhevol.2022.103250

Luncz, L. V., Mundry, R., & Boesch, C. (2012). Evidence for cultural differences between neighboring chimpanzee communities. Current Biology, 22(10), 922–926. 10.1016/j.cub.2012.03.031

Luncz, L. V., Slania, N. E., Almeida-Warren, K., Carvalho, S., Falótico, T., Malaivijitnond, S., Arroyo, A., de la Torre, I., & Proffitt, T. (2024). Tool skill impacts the archaeological evidence across technological primates. Scientific Reports, 14(1), 1–11. 10.1038/s41598-024-67048-z

Luncz, L. V., Svensson, M. S., Haslam, M., Malaivijitnond, S., Proffitt, T., & Gumert, M. (2017). Technological response of wild macaques (Macaca fascicularis) to anthropogenic change. International Journal of Primatology, 38, 872–880. 10.1007/s10764-017-9985-6

Malaivijitnond, S., Lekprayoon, C., Tandavanittj, N., Panha, S., Cheewatham, C., & Hamada, Y. (2007). Stone-tool usage by Thai long-tailed macaques (Macaca fascicularis). American Journal of Primatology, 69, 227–233. 10.1002/ajp.20342

Matuschek, H., Kliegl, R., Vasishth, S., Baayen, H., & Bates, D. (2017). Balancing Type I error and power in linear mixed models. Journal of Memory and Language, 94, 305–315. 10.1016/j.jml.2017.01.001

McGrew, W. C., Ham, R. M., White, L. J. T., Tutin, C. E. G., & Fernandez, M. (1997). Why don’t chimpanzees in Gabon crack nuts? International Journal of Primatology, 18(3), 353–374. 10.1023/A:1026382316131

Monteza-Moreno, C. M., Crofoot, M. C., Grote, M. N., & Jansen, P. A. (2020). Increased terrestriality in a Neotropical primate living on islands with reduced predation risk. Journal of Human Evolution, 143(102768). 10.1016/j.jhevol.2020.102768

Neadle, D., Bandini, E., & Tennie, C. (2020). Testing the individual and social learning abilities of task-naïve captive chimpanzees (Pan troglodytes sp.) in a nut-cracking task. PeerJ, 2020(3), 1–29. 10.7717/peerj.8734

Proffitt, T., Luncz, V. L., Malaivijitnond, S., Gumert, M., Svensson, M. S., & Haslam, M. (2018). Analysis of wild macaque stone tools used to crack oil palm nuts. Royal Society Open Science, 5(3). 10.1098/rsos.171904

Proffitt, T., Reeves, J. S., Pacome, S. S., & Luncz, L. V. (2022). Identifying functional and regional differences in chimpanzee stone tool technology. Royal Society Open Science, 9, 220826. 10.1098/rsos.220826

R Core Team. (2018). R: A Language and Environment for Statistical Computing. R Foundation for Statistical Computing. https://www.r-project.org/

Reeves, J. S., Proffitt, T., Almeida-Warren, K., & Luncz, L. V. (2023a). Modeling Oldowan tool transport from a primate perspective. Journal of Human Evolution, 181. 10.1016/j.jhevol.2023.103399

Reeves, J. S., Proffitt, T., & Luncz, L. V. (2021). Modeling a primate technological niche. Scientific Reports, 11, 23139. 10.1038/s41598-021-01849-4

Reeves, J. S., Proffitt, T., Malaivijitnond, S., & Luncz, L. V. (2023b). Emergent technological variation in archaeological landscapes: A primate perspective. Journal of the Royal Society Interface, 20(203). 10.1098/rsif.2023.0118

Reeves, J. S., Proffitt, T., Pacome, S. S., & Luncz, L. V. (2024). Searching for the earliest archaeological record: insights from chimpanzee material landscapes. Journal of the Royal Society Interface, 21, 20240101. 10.1098/rsif.2024.0101

Rutz, C., & St Clair, J. J. H. (2012). The evolutionary origins and ecological context of tool use in New Caledonian crows. Behavioural Processes, 89, 153–165. 10.1016/j.beproc.2011.11.005

Sanz, C. M., & Morgan, D. B. (2013). Ecological and social correlates of chimpanzee tool use. Philosophical Transactions of the Royal Society B, 368, 20120416. 10.1098/rstb.2012.0416

Schunk, L. (2021). Understanding Middle Palaeolithic asymmetric stone tool design and use: use-wear analysis and controlled experiments to assess Neanderthal technology. Johannes Gutenberg University Mainz.

Sirianni, G., Mundry, R., & Boesch, C. (2015). When to choose which tool: Multidimensional and conditional selection of nut-cracking hammers in wild chimpanzees. Animal Behaviour, 100, 152–165. 10.1016/j.anbehav.2014.11.022

Spagnoletti, N., Visalberghi, E., Verderane, M. P., Ottoni, E., Izar, P., & Fragaszy, D. (2012). Stone tool use in wild bearded capuchin monkeys, Cebus libidinosus. Is it a strategy to overcome food scarcity? Animal Behaviour, 83(5), 1285–1294. 10.1016/j.anbehav.2012.03.002

Van Schaik, C. P., Deaner, R. O., & Merrill, M. Y. (1999). The conditions for tool use in primates: Implications for the evolution of material culture. Journal of Human Evolution, 36(6), 719–741. 10.1006/jhev.1999.0304

Visalberghi, E., Addessi, E., Truppa, V., Spagnoletti, N., Ottoni, E., Izar, P., & Fragaszy, D. M. (2009a). Selection of effective stone tools by wild bearded capuchin monkeys. Current Biology, 19(3), 213–217. 10.1016/j.cub.2008.11.064

Visalberghi, E., Fragaszy, D. M., Ottoni, E., Izar, P., Oliveira M. G. de, & Andrade, F. R. D. (2007). Characteristics of hammer stones and anvils used by wild bearded capuchin monkeys (Cebus libidinosus) to crack open palm nuts. American Journal of Physical Anthropology, 132, 426–444. 10.1002/ajpa.20546

Visalberghi, E., Sabbatini, G., Taylor, A. H., & Hunt, G. R. (2017). Cognitive insights from tool use in nonhuman animals. In J. Call, G. M. Burghardt, I. M. Pepperberg, C. T. Snowdon, & T. Zentall (Eds.), APA handbook of comparative psychology: Perception, learning, and cognition (pp. 673–701). American Psychological Association. 10.1037/0000012-030

Visalberghi, E., Sirianni, G., Fragaszy, D. M., & Boesch, C. (2015). Percussive tool use by Taï western chimpanzees and Fazenda Boa Vista bearded capuchin monkeys: A comparison. Philosophical Transactions of the Royal Society B: Biological Sciences, 370, 20140351. 10.1098/rstb.2014.0351

Visalberghi, E., Spagnoletti, N., Ramos da Silva, E. D., Andrade, F. R. D., Ottoni, E., Izar, P., & Fragaszy, D. M. (2009b). Distribution of potential suitable hammers and transport of hammer tools and nuts by wild capuchin monkeys. Primates, 50, 95–104. 10.1007/s10329-008-0127-9

Vogel, E. R. (2005). Rank differences in energy intake rates in white-faced capuchin monkeys, Cebus capucinus: The effects of contest competition. Behavioral Ecology and Sociobiology, 58(4), 333–344. 10.1007/s00265-005-0960-4

Yamakoshi, G. (1998). Dietary responses to fruit scarcity of wild chimpanzees at Bossou, Guinea: Possible implications for ecological importance of tool use. American Journal of Physical Anthropology, 106(3), 283–295. 10.1002/(SICI)1096-8644(199807)106:3<283::AID-AJPA2>3.0.CO;2-O

Zuberbühler, K., & Jenny, D. (2002). Leopard predation and primate evolution. Journal of Human Evolution, 43(6), 873–886. 10.1006/jhev.2002.0605

