## Supplementary Information for "Material matters: raw material influences stone tool performance in capuchin monkeys"

### Material matters: variation in performance and durability of Capuchin percussive stone tools

#### RAW MATERIALS

**Supplementary Table 1:** The raw material and source of tool sets, and the tool-using population whose range the material was obtained from.

| Tool Set | | Raw Material | Source Location | Source population | Tool Size |
|---|---|---|---|---|---|
| 1 | *hammer* | phonolite | Olduvai Gorge, Tanzania | hominin | 257.6, 304.5, 580.6 (g) |
| | *anvil* | quartzite | Olduvai Gorge, Tanzania | hominin | 16 x 15.5 x 5 (cm) |
| 2 | *hammer* | siltstone | Phang-Nga, Thailand | macaque | 334.3, 448.1, 653.2 (g) |
| | *anvil* | siltstone | Phang-Nga, Thailand | macaque | 23 x 16 x 6 (cm) |
| 3 | *hammer* | siltstone | Phang-Nga, Thailand | macaque | 334.3, 448.1, 653.2 (g) |
| | *anvil* | concrete | Tietê, Brazil | capuchin | 27 x 20 x 2.5 (cm) |
| 4 | *hammer* | siltstone | Phang-Nga, Thailand | macaque | 334.3, 448.1, 653.2 (g) |
| | *anvil* | concrete | Tietê, Brazil | capuchin | 26 x 26 x 5 (cm) |
| 5 | *hammer* | quartzite | São Paulo, Brazil | capuchin | 317.8, 402.5, 625.2 (g) |
| | *anvil* | sandstone | São Paulo, Brazil | capuchin | 31 x 17.5 x 5.5 (cm) |
| 6 | *hammer* | quartzite | São Paulo, Brazil | capuchin | 317.8, 402.5, 312† (g) |
| | *anvil* | ironstone | Tietê, Brazil | capuchin | 19.5 x 15.5 x 6 (cm) |

† The large (625.2 g) quartzite hammerstone fractured into two c. 312 g pieces shortly after it was first used on the ironstone anvil.

Photographs of damaged raw material is provided, including anvils at the start and end of the study, and any fractured hammerstones.

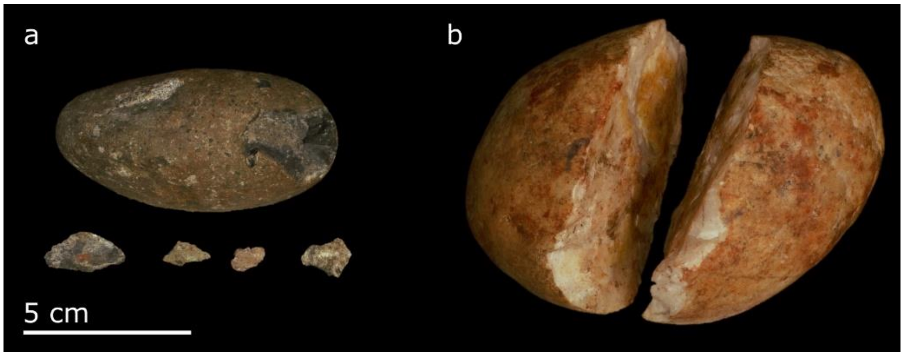


**Supplementary Figure 1: Hammerstone durability.** Damage to two of the hammerstones used in this study. The smallest phonolite hammer, sourced from the Oduvai Gorge in Tanzania, had some small fragments fall off (a). The largest quartzite hammer, sourced from Serra da Capivara National Park in São Paulo, Brazil, split in half (b).

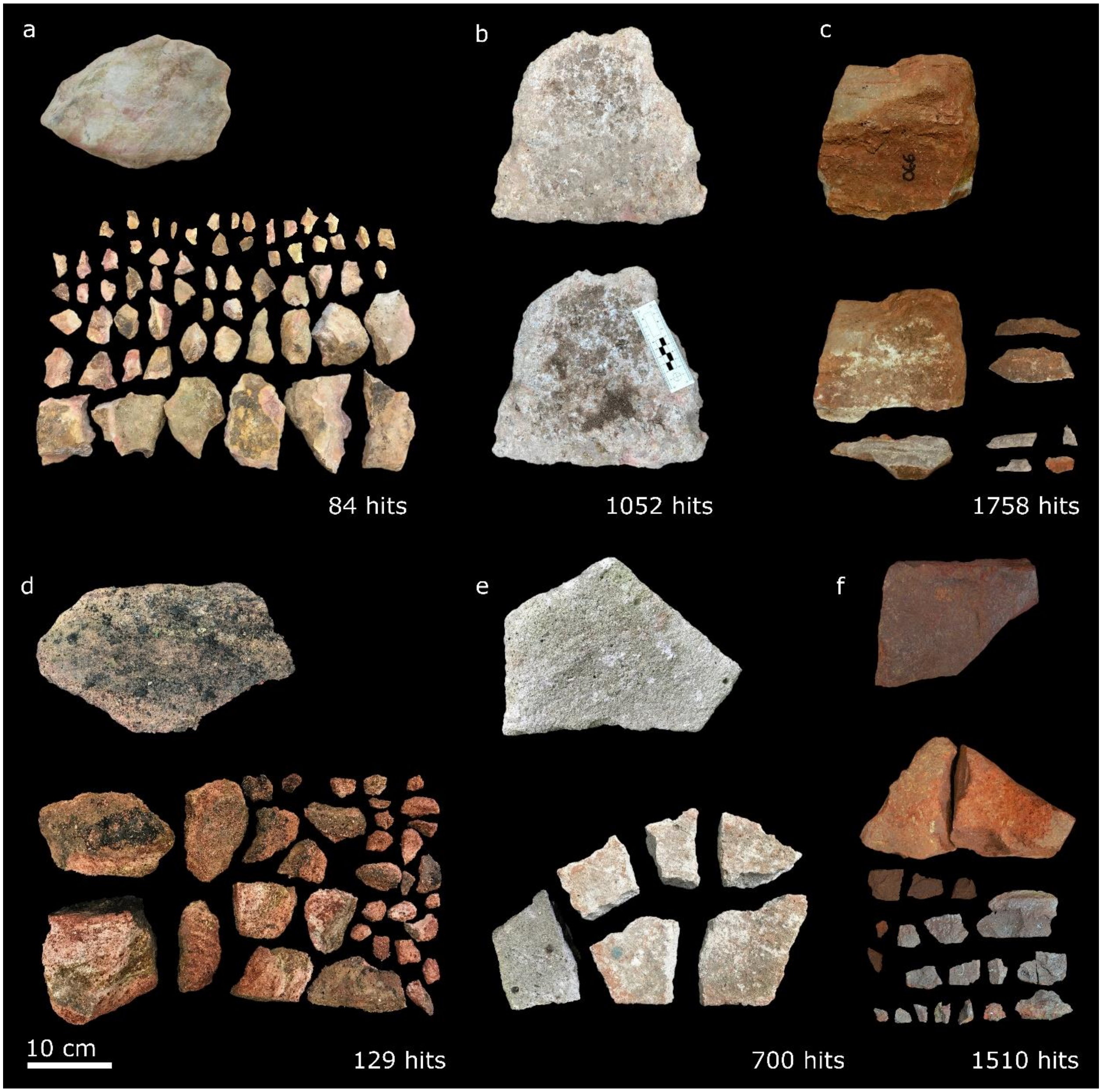


**Supplementary Figure 2: Anvil durability.** Photos of the (a) siltstone, (b) thicker concrete, (c) quartzite, (d) sandstone, (e) thinner concrete, and (f) ironstone anvils at the start (upper image) and end (lower image) of the study, alongside the total number of hammer stone hits sustained by the anvil. Photos exclude small (< 2cm) fragments.

#### MODEL DIAGNOSTICS

*Cracking Success*

The model was checked for complete separation for males and females, each tool set, and each individual. The only instance of complete separation was a female with five attempts at cracking a nut, none of which succeeded. Overdispersion was assessed using the DHARMa function 'testDispersion' (ref; Fig. 3a). This was deemed acceptable with a dispersion value close to 1 (1.02). Model stability was visually assessed by removing one individual at a time and rerunning the model. Results are summarized in Figure 3b. Stability was considered to be acceptable: the only parameter with a high level of instability was the effect of sex. The effect size for sex could vary widely if certain females were removed, but this was interpreted primarily as being a consequence of having a small number of females. This was considered acceptable as sex was not a variable of interest, and parameter estimates were stable for all other variables.

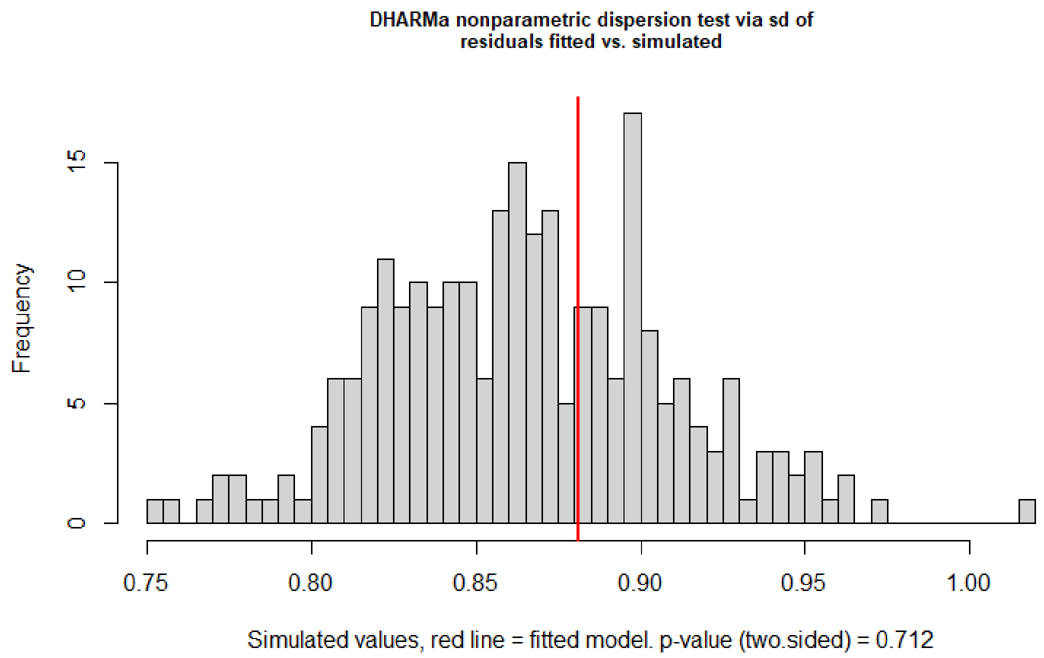


**Supplementary Figure 3a: Dispersion of the cracking success GLMM.** Plot shows distribution of fitted residuals against simulated, generated using DHARMa function 'testDispersion'.

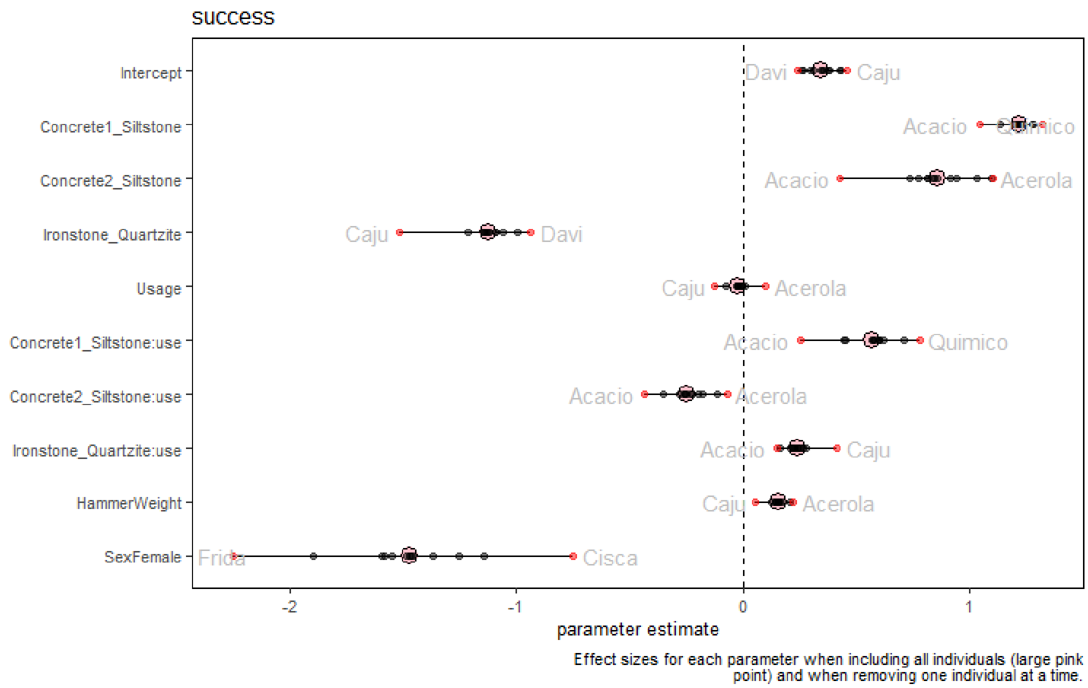


**Supplementary Figure 3b: Stability of the cracking success GLMM.** Parameter estimates with each individual removed. The original estimates are in pink. Each individual is represented by a black dot; red if it is the most extreme value for that parameter, along with that individual's name.

*Hits-per-nut*

The model was initially strongly right-skewed when using a zero-truncated Poisson distribution. This was changed to a zero-truncated negative binomial. With the negative binomial distribution the skew was much weaker and the model was not at all over dispersed (dispersion = 0.90; Fig 4a). Model stability was assessed by removing one individual at a time. The model was relatively stable (Fig. 4b), except for estimates regarding tool set 3 (Concrete1_Siltstone). If the individual Acerola was removed, then the estimate for this material increased, indicating more hits will be required for this tool set than if the individual was included. However, this was for the full model, which assumed that tool sets differed in how their performance changed over time (i.e. an interaction between material and usage). This interaction was not favoured; model comparison instead favoured the reduced model which had no such interaction. This was the case regardless of whether Acerola was included (full vs reduced model comparison: $X^2_3 = 5.2$, $p = 0.16$) or excluded ($X^2_3 = 7.0$, $p = 0.072$). Study interpretations are taken from the reduced model, which Acerola has minimal effect on (Fig. 4c). This individual was retained in the study.

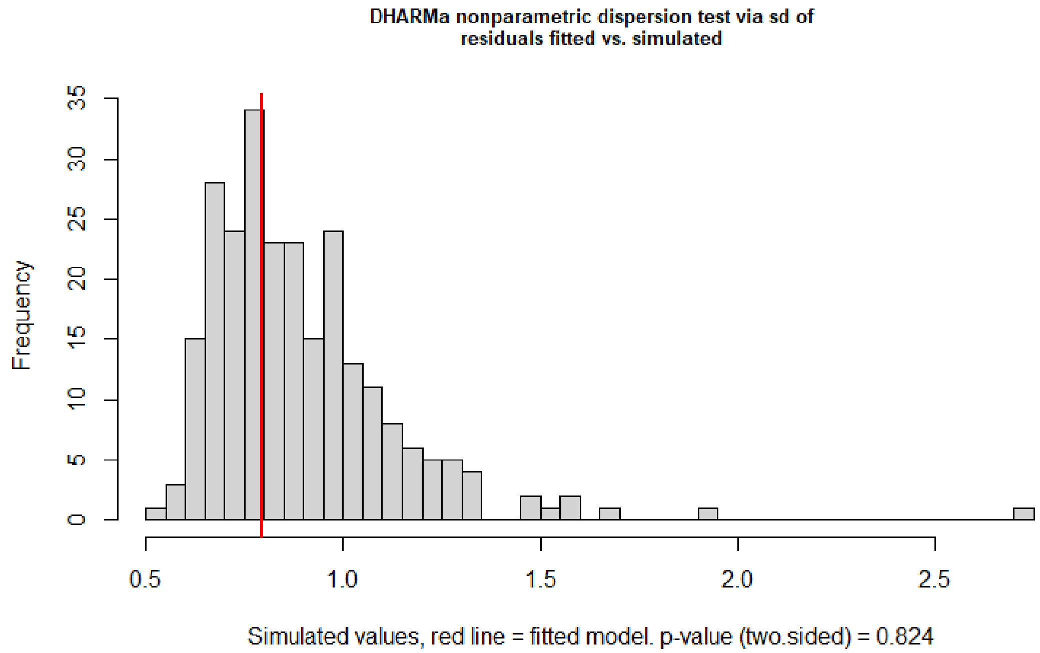


**Supplementary Figure 4a: Dispersion in the hits-per-nut GLMM. Plot** shows distribution of fitted residuals against simulated, generated using DHARMa function 'testDispersion'.

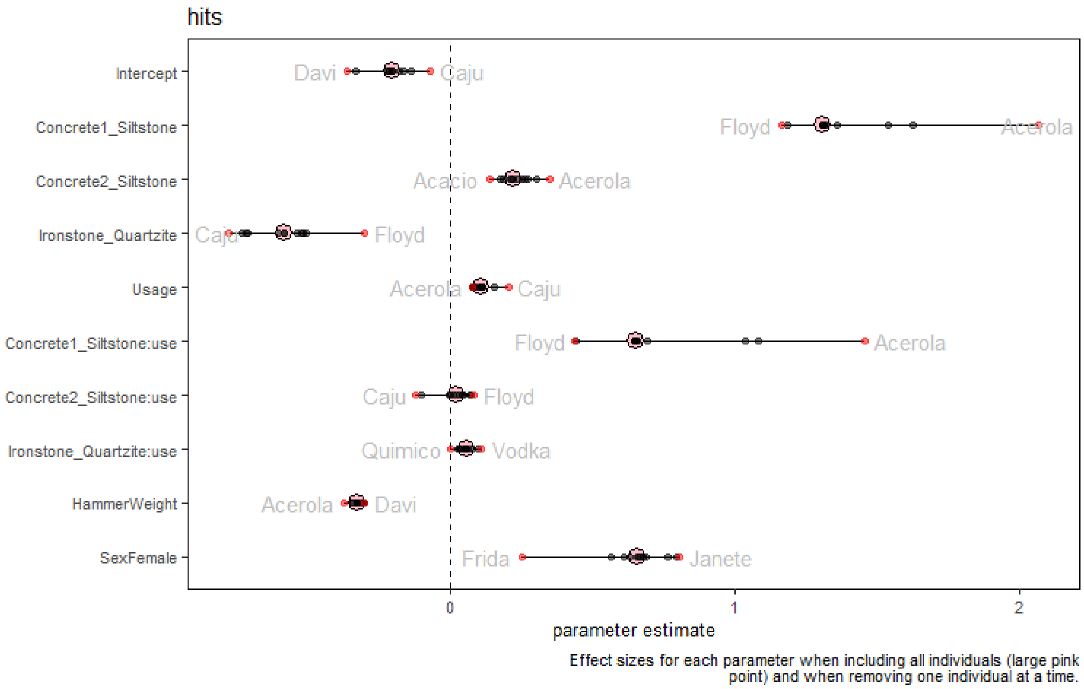


**Supplementary Figure 4b: Stability of the hits-per-nut GLMM.** Parameter estimates with each individual removed. The original estimates are in pink. Each individual is represented by a black dot; red if it is the most extreme value for that parameter, along with that individual's name.

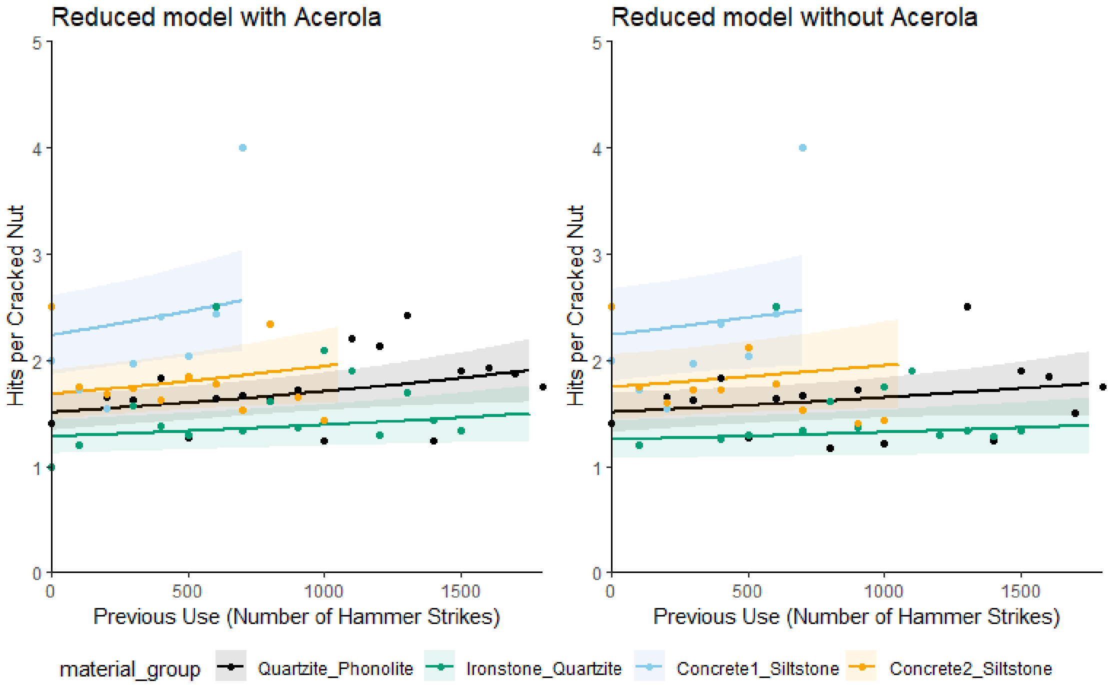


**Supplementary Figure 4c: Influence of the individual Acerola.** Plots show the predicted (lines) and observed (points) number of hits required to crack a nut when the individual Acacio is included (left) versus excluded (right) from the data. Results are for the reduced model, which is reported in the paper, where there is no interaction between material and use.

*Hits-per-food-reward*

This model was right skewed when using a zero-truncated Poisson distribution, but this skew was much less with a zero-truncated negative binomial distribution (Fig. 5a). Dispersion was also near 1 (dispersion = 0.92). Stability analysis, following the same protocol as for the other models, identified Acacio as being a potentially influential individual (Fig. 5b). Again, like for the hits-per-nut model, removing this individual made the values for tool set 3 (Concrete1_Siltstone) much larger (indicating many more hits required for this tool set). However, again, this was only for the full model where tool sets were allowed to vary in how they changed with use. Full versus null model reduction did not favour the inclusion of this interaction; this was true whether Acacio was included ($X^2_3 = 4.9$, $p = 0.18$) or excluded ($X^2_3 = 6.2$, $p = 0.10$) from the data. Acacio did not have a notable impact on the results of the reduced model where there is no interaction (Fig. 5c), which is the model from which this paper interpreted its results.

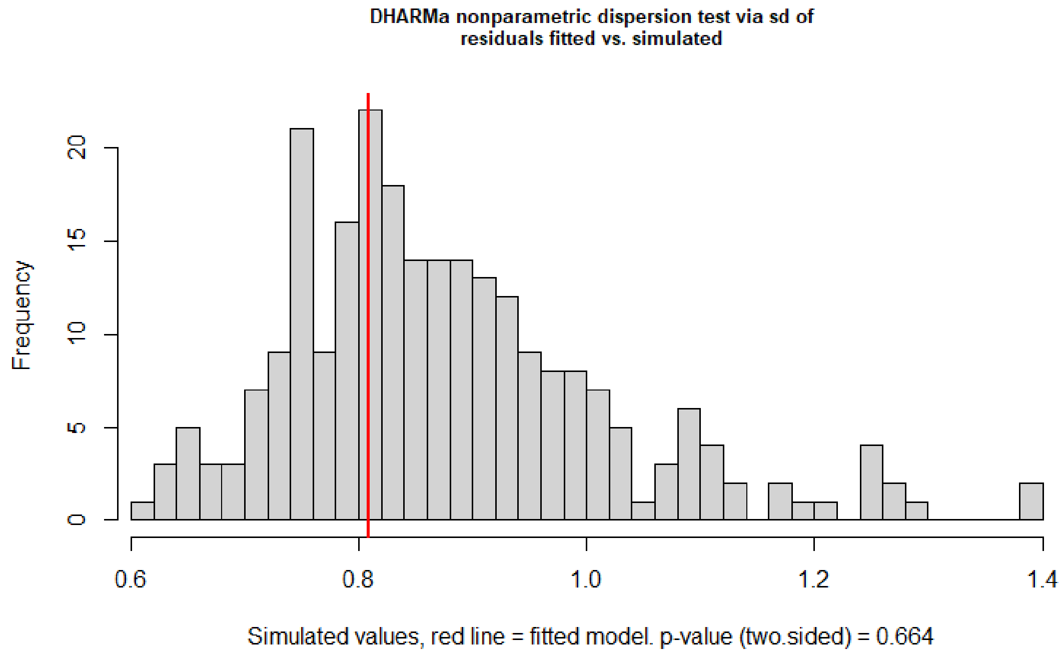


**Supplementary Figure 5a: Dispersion in the hits-per-reward GLMM.** Plot shows distribution of fitted residuals against simulated, generated using DHARMa function ‘testDispersion’.

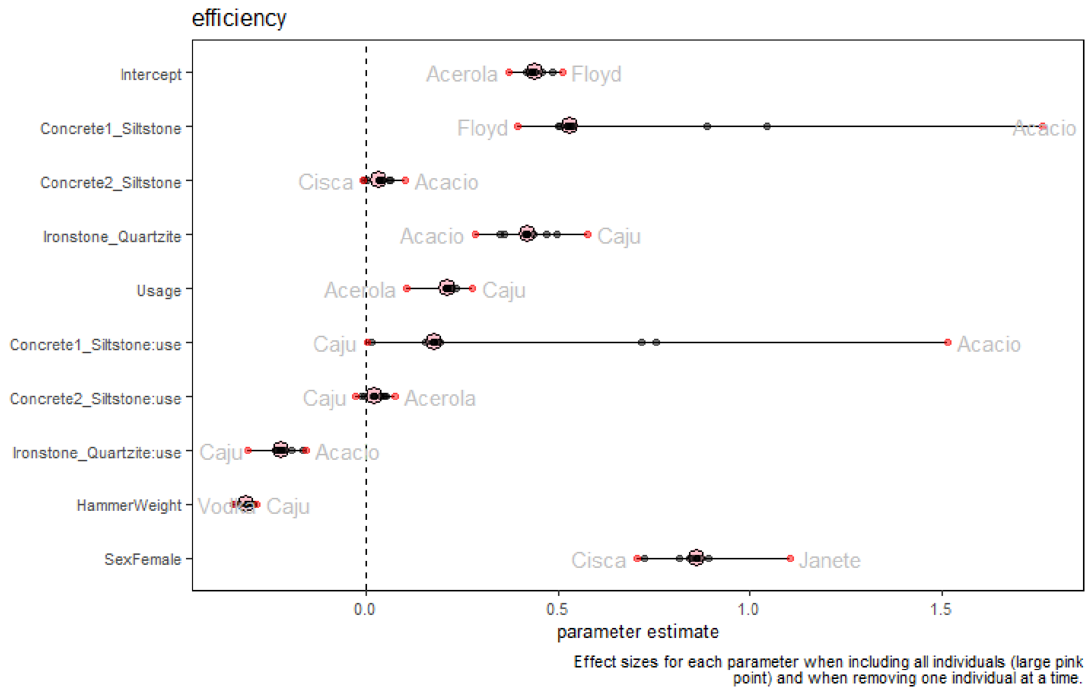


**Supplementary Figure 5b: Stability of the hits-per-reward GLMM.** Parameter estimates with each individual removed. The original estimates are in pink. Each individual is represented by a black dot; red if it is the most extreme value for that parameter, along with that individual’s name.

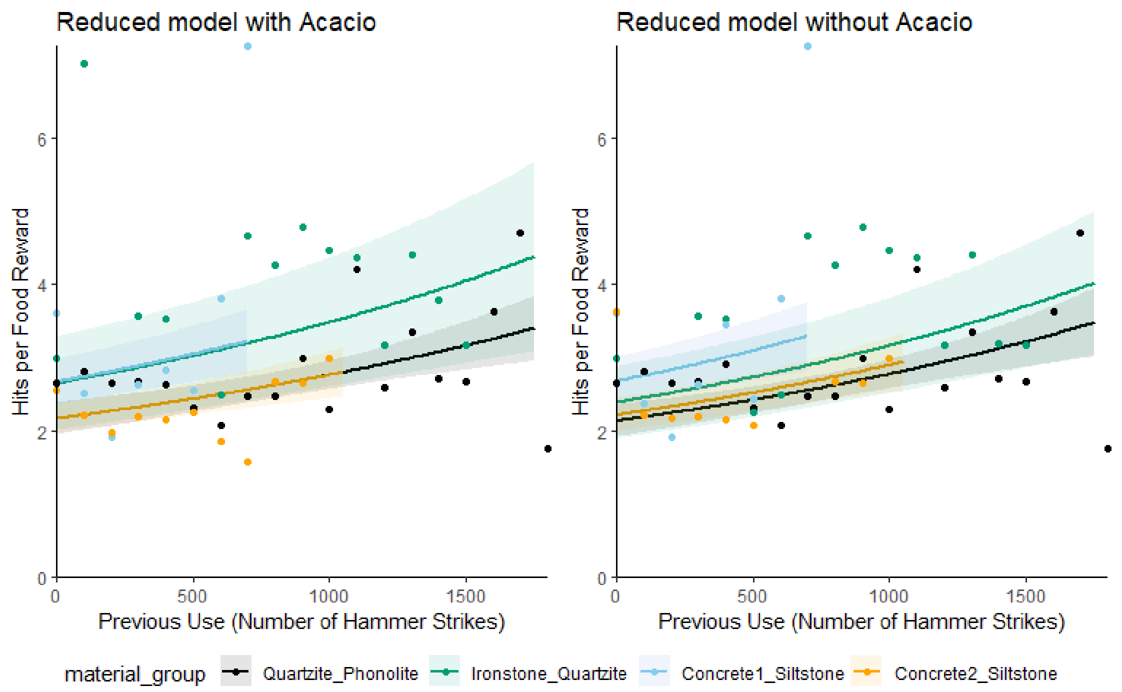


**Supplementary Figure 5c: Influence of the individual Acacio.** Plots show the predicted (lines) and observed (points) number of hits required to obtain a food reward when the individual Acacio is included (left) versus excluded (right) from the data. Results are for the reduced model, which is reported in the paper, where there is no interaction between material and use.

#### EXPANDED MODEL RESULTS

*Full versus reduced model comparisons*

**Supplementary Table 2:** Anova comparison of the full model (formula: response variable ~ material*use + hammer weight) against the null and reduced models. Three full models were tested with the following response variables: cracking success, hits-per-nut and hits-per-reward. Please note that all models also have a random effect of subject '(1 + material||subject)', and all models except the null have sex as a control variable.

| | | Success | | | Hits-Per-Nut | | | Hits-Per-Reward | | |
|---|---|---|---|---|---|---|---|---|---|---|
| Model | Model Formula | $X^2$ | *df* | *P* | $X^2$ | *df* | *P* | $X^2$ | *df* | *P* |
| Null | ~ 1 | 42.4 | 9 | < 0.001 | 65.8 | 9 | < 0.001 | 83.9 | 9 | < 0.001 |
| No Material | ~ use + hammer weight | 25.9 | 6 | < 0.001 | 25.5 | 6 | < 0.001 | 17.6 | 6 | < 0.01 |
| No Use | ~ material + hammer weight | 6.4 | 4 | 0.17 | 12.3 | 4 | <0.05 | 24.6 | 4 | < 0.001 |
| No Interaction | ~ material + use + hammer weight | 6.0 | 3 | 0.11 | 5.2 | 3 | 0.16 | 4.9 | 3 | 0.18 |
| No Hammer Weight | ~ material*use | 5.5 | 1 | <0.05 | 39.8 | 1 | < 0.001 | 40.7 | 1 | < 0.001 |

*Plotting Outliers*

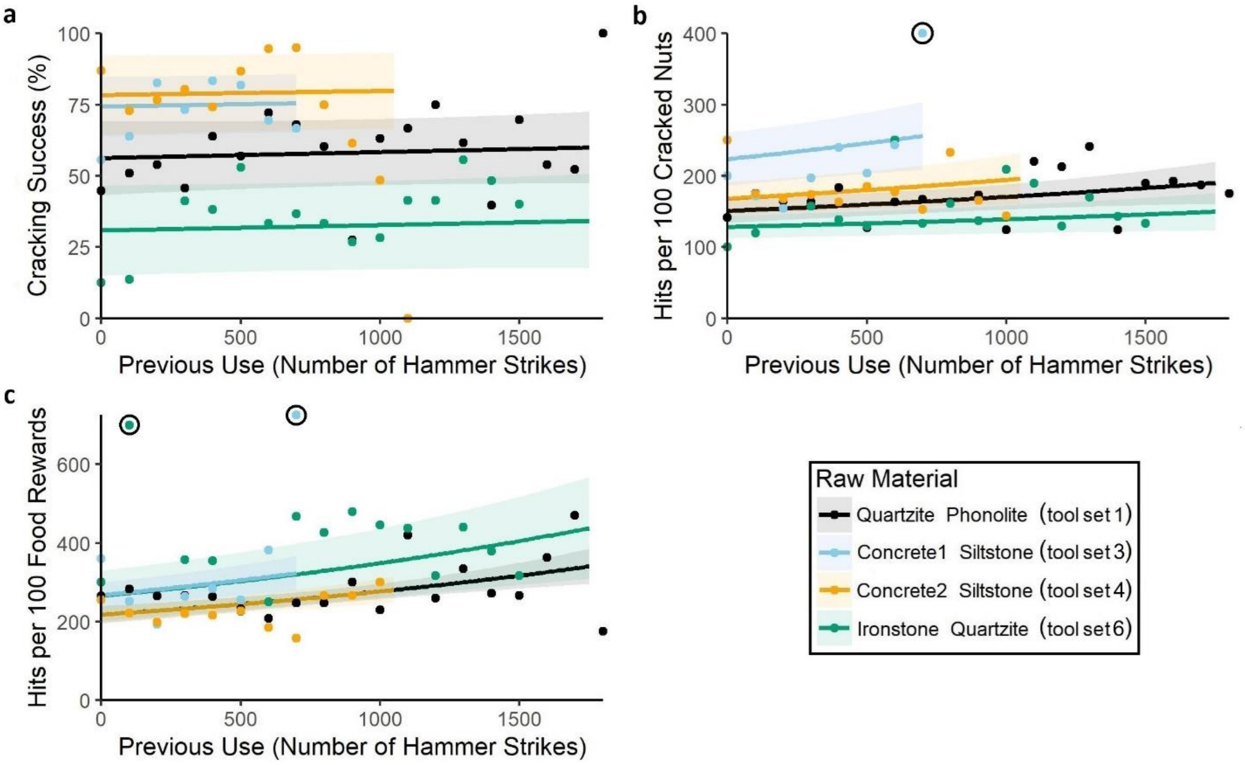


**Supplementary Figure 6: Outliers excluded from Figure 1 in the main paper.** *This figure is present in the main article as Figure 2, with three outlaying points excluded. The excluded points are included here and circled.* Tool performance as measured by the likelihood of successfully cracking a nut (a), hits to open a nut (b), and hits to obtain a food reward (c). Performance varies across tool sets of different raw material (listed as anvil then hammer stone) and with increasing use. Each point indicates the observed data, averaged across 100 hammer strikes. Lines reflect model predictions, assuming a male capuchin and hammer weight of 518 g, with no interaction between raw material and use. Shaded areas reflect one standard error either side of the estimate.

*Effect of hammer weight*

The primary aim of the paper was to investigate the influence of raw material and prior use on tool performance. However, we also found effects of hammer weight, as discussed in the full paper. The predicted effect of hammer weight is depicted in Supplementary Figure 6. This prediction assumes no interaction between raw material and past use of the tool, as such an interaction was not was not supported by full versus reduced model comparison for any of the three performance metrics ($P > 0.05$; SI Table 1).

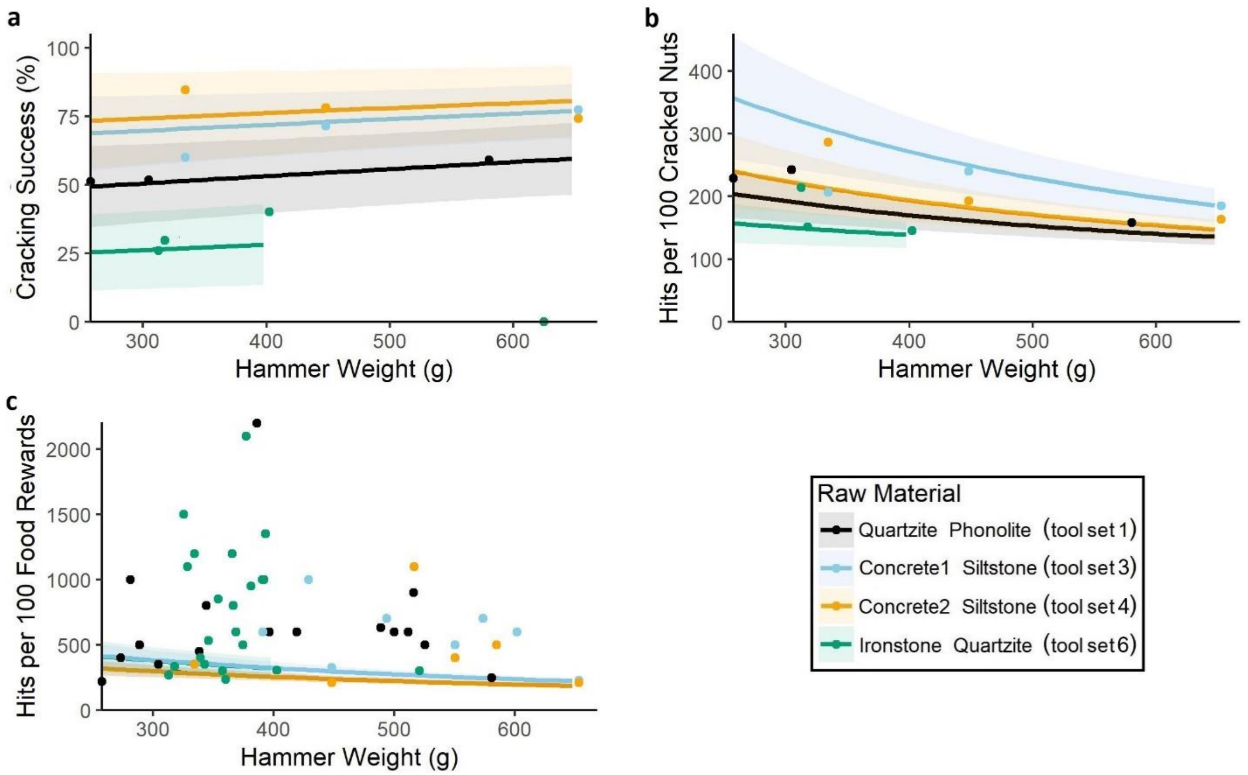


**Supplementary Figure 7: Influence of hammer weight on tool performance.** Tool performance as measured by the likelihood of successfully cracking a nut (a), hits to open a nut (b), and hits to obtain a food reward (c). Performance varies across tool sets of different raw material (listed as anvil then hammer stone) and with increasing hammer weight. Each point indicates the observed data, averaged for each hammer. When assessing the number of hits required for each food reward, sometimes multiple hammers were used. These weights were averaged. Lines reflect model predictions, assuming a male capuchin and hammer weight of 518 g, with the shaded area reflecting one standard error either side of the estimate.

*Full model results: allowing raw material and prior use to interact*

There was no evidence that raw materials differed in how their performance changed with prior use. For all three performance metrics (cracking success, hits per nut, hits per food reward) full versus reduced model comparison did not find a significant difference between the full model and one where the interaction between material and use was removed ($p > 0.05$). As such, results in the main text are described for the reduced version of each model, where there is no such interaction. Below we report the results of the full version of each model (Table 3, Fig. 8 and 9).

**Supplementary Table 3:** Results from the full cracking success, hits-per-nut and hits-per-reward models, where the interaction between material and use is included. Estimates for hammer weight and use have been scaled to reflect change per 100 grams and 100 cumulative hammer strikes respectively. The base material was tool set 1, which had a quartzite anvil and phonolite hammerstones (quartzite-phonolite). Material effects are from tool set 3 (concrete1-siltstone), tool set 4 (concrete2-siltstone) and tool set 6 (ironstone-quartzite).

| Parameter | Success | | | Hits-Per-Nut | | | Hits-Per-Reward | | |
|---|---|---|---|---|---|---|---|---|---|
| | Estimate | (SD) | *Z* | Estimate | (SD) | *Z* | Estimate | (SD) | *Z* |
| Intercept | 0.34 | (0.23) | 1.48 | -0.21 | (0.15) | -1.37 | 0.44 | (0.077) | 5.66 |
| Material | | | | | | | | | |
| *Tool Set 3* | 1.22 | (0.39) | 3.15 | 1.31 | (0.25) | 5.24 | 0.53 | (0.21) | 2.51 |
| *Tool Set 4* | 0.86 | (0.34) | 2.50 | 0.22 | (0.18) | 1.26 | 0.031 | (0.11) | 0.27 |
| *Tool Set 6* | -1.13 | (0.30) | -3.77 | -0.59 | (0.27) | -2.21 | 0.42 | (0.19) | 2.25 |
| Use | -0.0048 | (0.016) | -0.30 | 0.023 | (0.015) | 1.50 | 0.046 | (0.011) | 4.33 |
| Use*Material | | | | | | | | | |
| *Use*Tool Set 3* | 0.57 | (0.42) | 1.35 | 0.65 | (0.29) | 2.21 | 0.18 | (0.23) | 0.78 |
| *Use*Tool Set 4* | -0.25 | (0.25) | -0.99 | 0.02 | (0.17) | 0.11 | 0.021 | (0.12) | 0.17 |
| *Use*Tool Set 6* | 0.24 | (0.14) | 1.71 | 0.054 | (0.16) | 0.33 | -0.22 | (0.11) | -1.93 |
| Hammer Weight | 0.12 | (0.052) | 2.35 | -0.25 | (0.04) | -6.36 | -0.24 | (0.038) | -6.39 |
| Sex *Female* | -1.47 | (0.49) | -2.99 | 0.66 | (0.32) | 2.02 | 0.86 | (0.22) | 3.92 |

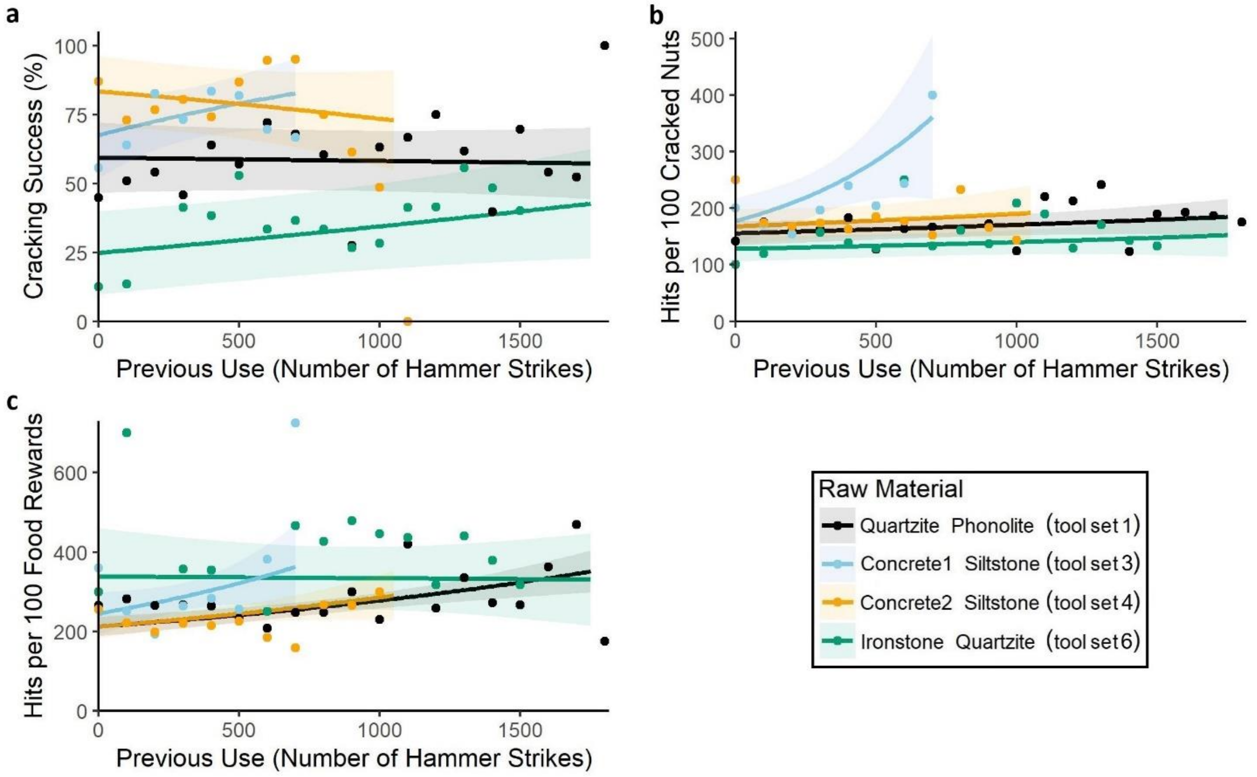


**Supplementary Figure 8: Influence of raw material and prior use on tool performance – full model results.** Tool performance from the full model for cracking success (a), hits to open a nut (b), and hits to obtain a food reward (c). In the full model, there is an interaction between material and use. Performance varies across tool sets of different raw material (listed as anvil then hammer stone) and with increasing use. Each point indicates the observed data, averaged across 100 hammer strikes. Lines reflect model predictions, assuming a male capuchin and hammer weight of 518 g, with the shaded area reflecting one standard error either side of the estimate.

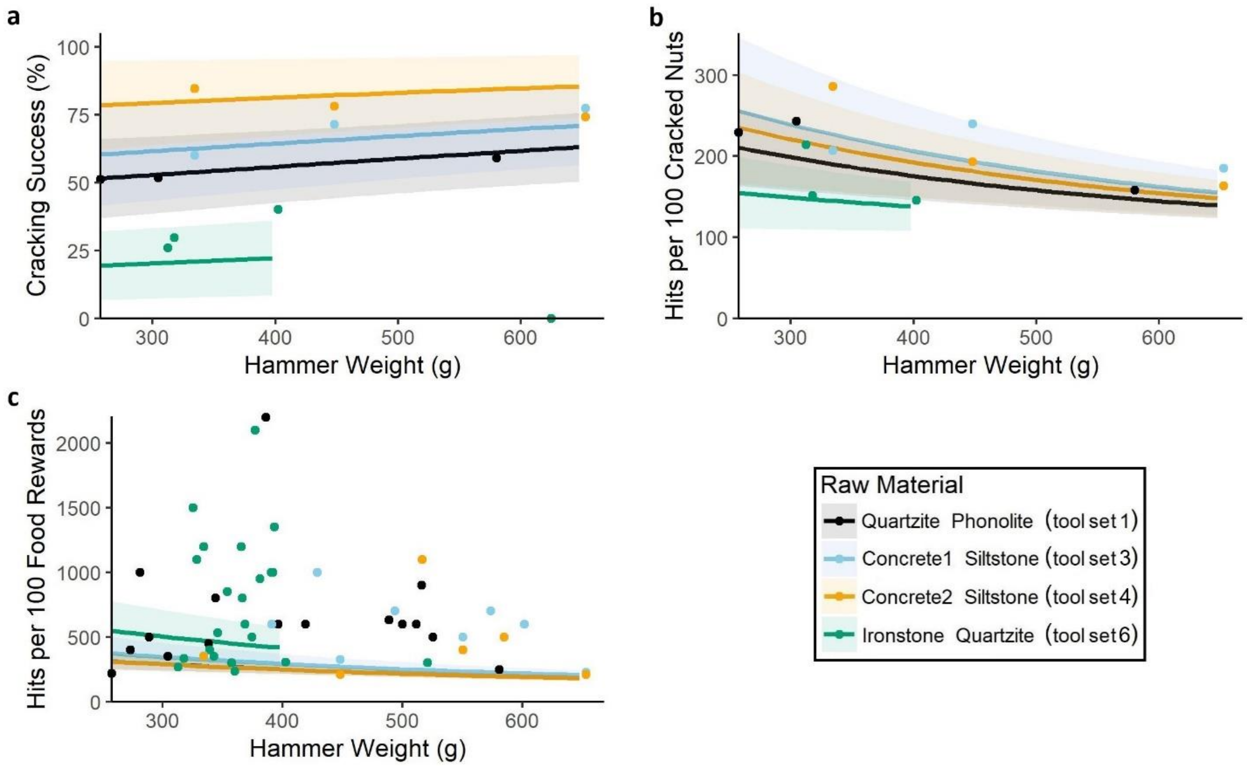


**Supplementary Figure 9: Influence of hammer weight on tool performance – full model results.** Tool performance from the full model for cracking success (a), hits to open a nut (b), and hits to obtain a food reward (c). In the full model, there is an interaction between material and use. Performance varies across tool sets of different raw material (listed as anvil then hammer stone) and with increasing hammer weight. Each point indicates the observed data, averaged for each hammer. When assessing the number of hits required for each food reward, sometimes multiple hammers were used. These weights were averaged. Lines reflect model predictions, assuming a male capuchin and hammer weight of 518 g, with the shaded area reflecting one standard error either side of the estimate.
